# Integrative Omics reveals genetic basis and TaMYB7-A1’s function in wheat WUE and drought resilience

**DOI:** 10.1101/2024.12.12.628287

**Authors:** Yuxin Zhou, Hao Wang, Yunzhou Qiao, Peng Zhao, Yuan Cao, Xuemei Liu, Yiman Yang, Xuelei Lin, Shengbao Xu, Baodi Dong, Dongzhi Wang, Jun Xiao

## Abstract

Improving water use efficiency (WUE) and drought resistance in wheat is critical for ensuring global food security under changing climate conditions. Here, we integrated multi-omic data, including population-scale phenotyping, transcriptomics, and genomics, to dissect the genetic and molecular mechanisms underlying WUE and drought resilience in wheat. Genome-wide association studies (GWAS) revealed 8,135 SNPs associated with WUE-related traits, identifying 258 conditional and non-conditional QTLs, many of which co-localized with known drought-resistance genes. Pan-transcriptome analysis uncovered tissue-specific expression patterns, core and unique gene functions, and dynamic sub-genomic biases in response to drought. eQTL mapping pinpointed 146,966 regulatory loci, including condition-specific hotspots enriched for genes involved in water regulation, osmoregulation, and photosynthesis. Integration of Weighted gene co-expression network analysis (WGCNA), Summary-data-based Mendelian Randomization (SMR) and GWAS, eQTLs identified 207 candidate causal genes as key regulators for WUE-related traits in wheat, such as TaMYB7-A1. Functional analyses found that TaMYB7-A1 enhances drought tolerance by promoting root growth, reducing oxidative stress, and improving osmotic regulation, enabling better water access and survival under stress. It also increases photosynthesis efficiency and WUE, boosting yield under drought without compromising performance in well-watered conditions, making it ideal target for breeding. Our findings provide a comprehensive omic framework for understanding the genetic architecture of WUE and drought resistance, offering valuable targets for breeding resilient wheat varieties.

## Introduction

Common wheat (*Triticum aestivum* L.) is a crucial global staple, providing around 20% of daily caloric intake and protein worldwide^1^. With a growing human population and intensifying climate change, the demand for wheat is increasing alongside the prevalence of water scarcity, particularly in arid and semi-arid regions where wheat production is already constrained by limited water availability^2,3^. Water deficiency affects approximately 40% of global cropland, reducing wheat yield potential (estimated to cause ∼30% yield losses) and severe impacts grain quality, thus posing a significant risk to global food security and nutrition^2,4^. Given that agriculture accounts for 70% of global water consumption, improving water use efficiency (WUE) in crops is essential for optimizing production under water-limited conditions^5^. Developing wheat varieties with enhanced drought resilience has become a research priority to ensure sustainable agricultural productivity and to secure future food supplies amid evolving climatic challenges^6^.

Drought stress profoundly affects wheat growth and development, leading to substantial yield losses^7,8^. To cope with drought, wheat activates various physiological and biochemical mechanisms to reduce water loss and enhance stress tolerance^9^. A key adaptation involves modifying root architecture, increasing root depth and biomass to optimize water uptake from deeper soil layers^10^. Additionally, drought induces the accumulation of abscisic acid (ABA), which promotes stomatal closure to minimize transpiration and conserve water^8^. However, this response limits COL uptake and photosynthesis, creating a trade-off between water conservation and carbon assimilation^11,12^. ABA signaling is crucial for activating drought-responsive genes such as NAM [No Apical Meristem]–ATAF [Arabidopsis Transcription Activation Factor]–CUC [Cup-shaped Cotyledons] transcription factor (TF) *TaNAC48*^13^, *Delta-1-Pyrroline-5-Carboxylate Synthase 1* (*TaP5CS1*)^14^, and DREB type TF *TaDTG6-B*^15^, orchestrating a complex network of drought responses. Manipulating key ABA signaling factors like *drought-induced wilting 1* (*DIW1*)^16^, ABA receptors PYRABACTIN RESISTANCE1 (PYR1)/PYR1-LIKE (PYL)/REGULATORY COMPONENTS OF ABA RECEPTORS (RCAR) gene *TaPYL1-1B*^17^, *TaPYL4*^18^ and *TaPYL9*^19^ have improved wheat’s capacity to survive under water-limiting conditions.

Drought also increases reactive oxygen species (ROS), which, while potentially damaging, act as signaling molecules to activate antioxidant defenses^20,21^. Wheat enhances its antioxidant enzyme systems, including superoxide dismutase (SOD) and catalase (CAT), to mitigate oxidative damage^22^. Moreover, osmoprotectants such as proline and soluble sugars accumulate within cells, stabilizing cellular structures and maintaining osmotic balance^23^. The calcium ion (Ca²L) signaling pathway is essential, as fluctuations in Ca²L levels activate drought-responsive genes and proteins, regulating WUE and adaptation to prolonged water scarcity^24^. These intricate signaling and biochemical adjustments enable wheat to survive and grow under drought conditions, though often with reduced yield potential^25^. While drought resistance mechanisms ensure survival under water deficit, WUE emphasizes maintaining high productivity with minimal water usage throughout growth stages^26,27^.

Advances in quantitative trait loci (QTL) mapping, genome wide association studies (GWAS), and mutation screening have helped uncover factors linked to drought tolerance and to less extend of WUE in wheat ^6,28,29^. Key TF families implicated in drought response include NAC (e.g., TaNAC071)^30^, AP2/ERF (e.g., TaDTG6)^15^, MYB/MYC (e.g., TaMYB30-B)^31^, bZIP (e.g., TabZIP156)^32^, and WRKY (e.g., TaWRKY31)^33^, which modulate stress-responsive genes. TabHLH27 regulates both drought responsive genes and developmental factors to balance root growth and drought tolerance, thus promoting WUE^34^. Additionally, other type of functional proteins target drought adaptation via various mechanisms: LATERAL ROOT DENSITY (LRD)^35^ and SINA-type E3 ubiquitin ligase TaSINA2B^36^ aid root growth, vacuolar cation/H+ antiporters *TaNHX2* and *TaNHX3* influence stomatal regulation^37^, alkane biosynthesis enzyme ECERIFERUM1-6A (TaCER1) promotes cuticular wax production^38^, and WD40 domain containing TaWD40-4B.1^39^ and VQ protein TaVQ4-D^40^ support ROS scavenging. Overall, the limited knowledge of the genetic control of drought resistance and WUE constrains molecular breeding efforts aimed at improving these traits in wheat^6,41^.

With genetic variants shown to modulate complex traits via gene expression, integrative analyses, including DNA variation, expression diversity, and transcriptional regulatory variability have been used to reveal the core network and module factors of growth or stress response. Recent studies in maize, soybean, cotton and wheat have successfully integrated genomic, transcriptomic, and epigenomic data, shedding light on regulatory networks that drive critical traits like cotton fiber development^42,43^, wheat spike architecture^44^, and seed biosynthesis processes (oil content, storage protein, starch)^45,46^, as well as responses to abiotic stress^47–49^. For example, Yuan et al. combined GWAS, expression quantitative trait loci (eQTLs), and transcriptome-wide association studies (TWAS) data from 421 soybean accessions to map the genetic architecture of seed weight and oil content^45^; Yang et al. performed GWAS and eQTL analysis of 107 accessions and identified 86 QTLs, 105,462 eQTLs, and 68 eQTL hotspots associating with drought tolerance in emmer wheat^49^. This integrated, systems-level approach has opened new avenues in plant biology, allowing researchers to pinpoint regulatory genes and pathways that single-omics studies often miss.

To unravel the molecular networks underlying drought resistance and WUE in wheat, we generated a genomic regulatory landscape of wheat seedlings across varying water conditions using genetic variation, transcriptome data, and WUE-related traits from 228 wheat accessions. By integrating GWAS, eQTL, Weighted gene co-expression network analysis (WGCNA), and Summary-data-based Mendelian Randomization (SMR) analyses, we identified and prioritized causal genes critical for wheat’s adaptation to water deficit, via a small-scale validation using KN9204 gene-indexed mutant population^50^. Our study highlights the R2R3-MYB transcription factor, TaMYB7-A1, which enhances drought tolerance by promoting higher WUE and stable osmotic adjustment. This integrated approach provides a valuable resource for understanding wheat’s response to water stress at both the population and genome-wide levels, offering new perspectives for pinpointing essential genes and genetic loci.

## Results

### Genetic dissection of water use efficiency in common wheat through genome-wide association studies

Selecting appropriate drought treatment conditions is critical for accurately capturing phenotypic differences among wheat genotypes under varying environments^51^. To determine optimal conditions for assessing drought vulnerability in wheat varieties, we measured biomass and water use efficiency (WUE_p_, defined as dry biomass per unit water cost) at the seedling stage across a gradient of Relative Soil Water Content (RSWC) conditions. This assessment included wheat varieties previously categorized as drought-sensitive, drought-tolerant, and moderately drought-tolerant^52^. Overall, biomass increased with higher RSWC levels, with drought-sensitive varieties showing significantly reduced biomass at 6% RSWC, while drought-tolerant varieties maintained relatively high biomass at 15% RSWC. In contrast, WUE_p_ displayed an inverse trend, decreasing as RSWC increased, forming an S-shaped curve with an inflection point at 9% RSWC **(Fig. 1a**). Based on these findings, RSWC values of 15%, 9%, and 6% were designated as well-watered (WW), moderate deficit soil water (DS1), and severe deficit soil water (DS2) conditions, respectively.

**Fig. 1.**
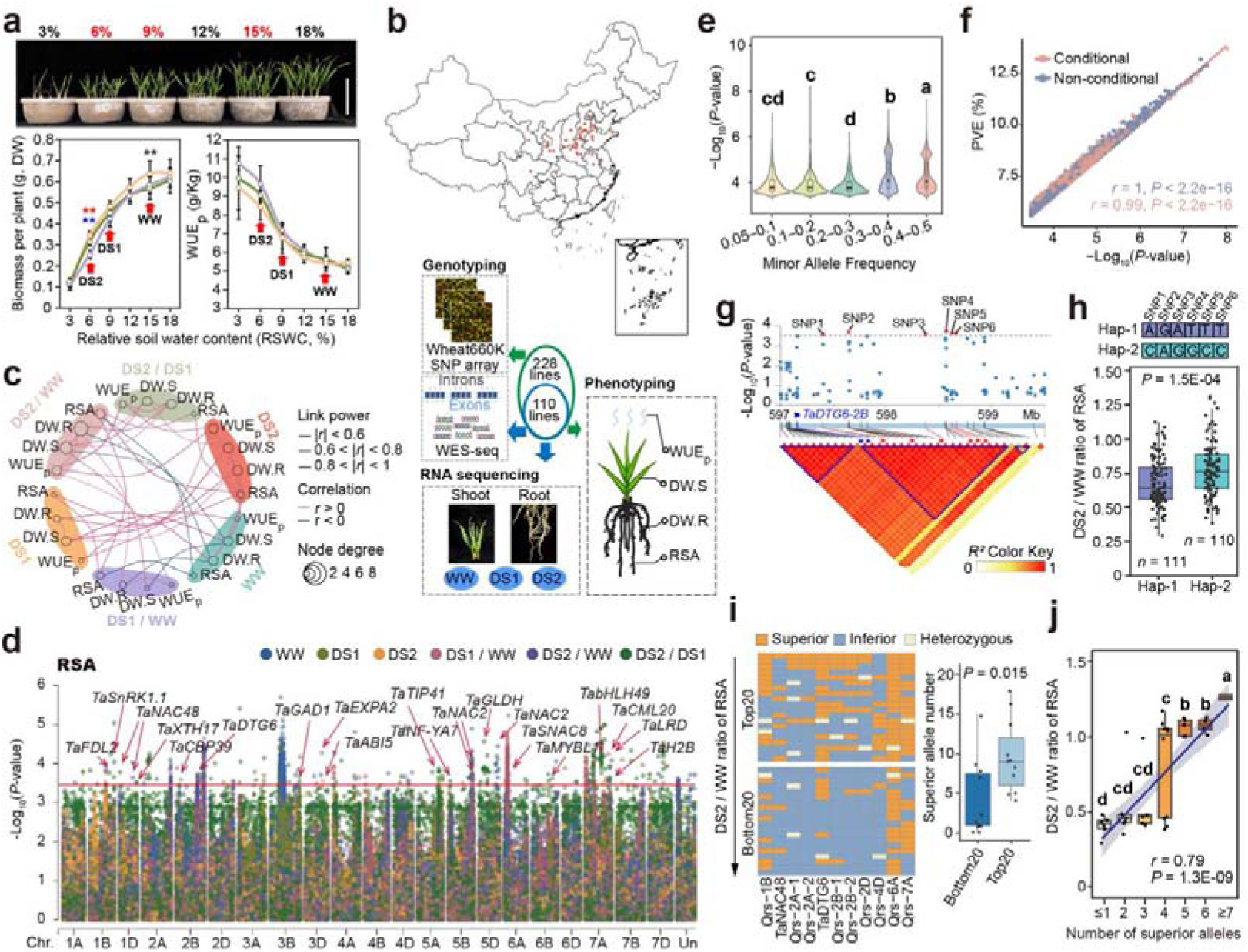
GWAS analysis identifying genetic variations contributing to the water use efficiency (WUE) in wheat. **(a)** Illustration showing the typical appearance of seedlings under varying relative soil water content (RSWC) conditions, alongside biomass-RSWC and WUE_p_-RSWC curves for drought-sensitive, drought-tolerant, and drought-moderately-tolerant wheat varieties. The RSWC level used to assess drought in wheat population were indicated by a red arrow. Error bars represent the standard deviation (S.D.) calculated from the five varieties in each group. Statistical significance of differences in biomass and WUE_p_ under each RSWC condition was determined using Student’s *t*-test. Significant differences are marked by the corresponding colors: red, drought-sensitive varieties vs. drought-tolerant varieties; blue, drought-sensitive varieties vs. drought-moderately-tolerant varieties; black, drought-moderately-tolerant varieties vs. drought-tolerant varieties. *, *P* ≤ 0.05; **, *P* ≤ 0.01; ns, no significant difference. Scale bar = 10 cm. DW, dry weight. **(b)** Geographic distribution, phenotyping, genotyping, and RNA-seq experimental design for the accessions. Each accession is displayed as a red dot. The 228 accessions were used for phenotyping and genotyped using Wheat660K SNP array, the 110 core set were performed to Whole Exome Sequencing (WES-seq) and RNA-seq. **(c)** Correlation Networks Across Different Traits. The traits under each condition were clustered with the same background color. Only edges with an average |*r*| > 0.6 are displayed. The correlation strength is indicated by the thickness of the edges, with positive correlations shown in pink and negative correlations in blue. The number of correlations with |*r*| > 0.6 is represented by the size of the nodes. **(d)** Manhattan plots illustrating the Llog_10_(*P*-value) values for root surface area (RSA) across multiple environments. Horizontal line indicates the genome-wide significant threshold of Llog_10_(*P*-value) = 3.50. Dots of different colors represent different environments. Wheat orthologs of reported drought-resistance genes located in the linkage disequilibrium (LD) region were labeled. **(e)** Comparison of the Llog_10_(*P*-value) values in GWAS for SNPs with different minor allele frequency. SNPs passed the significance threshold of Llog_10_(*P*-value) = 3.50 were retained, and data from different traits and conditions were polled together. Tukey’s HSD multiple comparison test was used to compare the Llog_10_(*P*-value) value between each group, with different letters indicate a statistically significant difference at *P* < 0.05. **(f)** The scatterplots showing the correlation between the Llog_10_(*P*) value and marker *R^2^*value of SNPs in GWAS. SNPs passed the significance threshold of Llog_10_(*P*-value) = 3.50 were retained, and data from different traits and conditions were polled together. The *r* value and *P* values were determined by Pearson correlation coefficient analysis. **(g)** Local Manhattan plot and LD heatmap focus on single nucleotide polymorphisms (SNPs) centre around the GWAS signal of *qRSA.DS2/WW-2B*. In the Manhattan plot, SNPs significantly associated with RSA.DS2/WW are marked with red dots, indicated by red asterisks in the LD heatmap, and connected by red lines. *TaDTG6-2B* is highlighted in blue, with SNPs in its genomic region represented by blue asterisks in the LD heatmap and connected by blue lines. The color intensity from white to red represents *r²* values ranging from 0 to 1. **(h)** Box plot comparing RSA.DS2/WW between haplotypes of *qRSA.DS2/WW-2B*. Wilcoxon rankLsum test was used to determine the statistical significance between two groups. Borders represent the first and third quartiles, center line denotes median, and whiskers extend to 1.5 times the interquartile range beyond the quartiles. **(i)** The aggregation of superior alleles in the top and bottom 20 accessions with highest and lowest RSA.DS2/WW content in the GWAS panel. Wilcoxon rankLsum test was used to determine the statistical significance between two groups. In the box plot, borders represent the first and third quartiles, center line denotes median, and whiskers extend to 1.5 times the interquartile range beyond the quartiles. **(j)** The scatterplots showing RSA.DS2/WW values plotted against the accumulation of superior alleles. The blue line represents the linear trend, with the grey shade indicating the 95% confidence interval of the trend. The *r* value and *P* values were determined by Pearson correlation coefficient analysis. Tukey’s multiple comparisons test was applied to assess the significance of RSA.DS2/WW among accessions carrying different numbers of superior alleles, with different letters indicating a significant difference at *P* < 0.05. In the box plot, the borders represent the first and third quartiles, the center line denotes the median, and whiskers extend to 1.5 times the interquartile range beyond the quartiles.

We then evaluated a diverse population of 228 wheat accessions, representing a broad spectrum of great genetic diversity and geographical distribution of wheat in China^34^, Each accession was assessed for WUE_p_, root dry weight (DW.R), shoot dry weight (DW.S), and root surface area (RSA) at the seedling stage under WW, DS1, and DS2 conditions **(Fig. 1b and Supplementary Table 1**). The 228 accessions exhibited significant variation across all measured traits, with DW.R, DW.S, and WUE_p_ strongly correlated, while RSA showed weaker correlations with other traits **(Fig. 1c and Supplementary Fig. 1a**). On average, DW.R, DW.S, and RSA progressively decreased as drought severity increased from WW to DS1 and DS2, but individual accessions exhibited distinct drought sensitivities (**Supplementary Fig. 1b, c**). Notably, WUE_p_ demonstrated greater variability under DS1 and DS2 conditions compared to WW. These extensive phenotypic variations provide a rich resource for uncovering genetic factors associated with drought-resistance and WUE_p_, and valuable germplasms for breeding programs aimed at enhancing drought resilience and optimizing WUE in wheat (**Supplementary Fig. 1c**).

The 228 wheat accessions were genotyped using the Wheat660K single-nucleotide polymorphism (SNP) array^53^, yielding 323,741 high-quality SNPs after stringent filtering (see methods for details). This comprehensive dataset enabled the classification of the diverse panel into three distinct sub-populations (**Supplementary Fig. 2a**). The analysis revealed an average linkage disequilibrium (LD) decay distance of 3 Mb (**Supplementary Fig. 2b**), aligning with previous reports^54,55^. To dissect the genetic basis of variations in drought vulnerability and WUE, we conducted a Genome-Wide Association Study (GWAS) using a mixed linear model (MLM) that accounts for population structure and kinship. This analysis identified 8,135 unique SNPs significantly associated with RSA, DW.R, DW.S, and WUE_p_ traits **(Fig. 1d, and Supplementary Fig. 2c, d**). In total, we identified 141 non-conditional QTLs (associated with absolute measured trait values) and 117 conditional QTLs (associated with trait ratios between different water conditions) **(Fig. 1d, Supplementary Fig. 2d-f, and Supplementary Table 2**). Interestingly, SNPs with larger minor allele frequencies (MAF) exhibited higher detection power and stronger allele effects **(Fig. 1e and Supplementary Fig. 2g**). Moreover, a significant positive correlation was observed between SNP association strength and its contribution to phenotypic variation, both for conditional and non-conditional QTLs **(Fig. 1f**).

Notably, many wheat orthologs of known drought-resistance genes co-localized with our GWAS signals **(Fig. 1d and Supplementary Fig. 2d**). For instance, *TaDTG6-B*, known to enhance drought tolerance in wheat^15^, was located within the LD block of *qRSA.DS2/WW-2B*, approximately 1.56 Mb from the leading SNP *AX-110493888* **(Fig. 1g**). Accessions carrying the Hap2 allele (*n* = 110) of significant SNPs associated with *AX-110493888* exhibited significantly higher RSA.DS2/WW values compared to those with the Hap1 allele (*n* = 111) **(Fig. 1h**). To further evaluate the contributions of superior alleles across multiple QTL loci to RSA.DS2/WW, we selected the top and bottom 20 accessions of RSA values from the GWAS panel and analyzed the aggregation of superior alleles. A significant difference in the number of superior alleles was observed between these groups **(Fig. 1i**). Accessions with higher RSA.DS2/WW values contained a greater number of superior alleles compared to those with lower values **(Fig. 1j**). This suggests that superior alleles function additively to enhance RSA in wheat under drought conditions.

Thus, we developed a system to analyze seedling-stage water use efficiency in wheat, identifying QTLs linked to RSA, DW.S, DW.R, and WUE_p_, including known and novel drought-resistance factors.

### Pan-transcriptome analysis reveals gene expression dynamics and sub-genome bias under drought

Gene expression variation is a major driver of phenotypic diversity, making the pan-transcriptome an essential tool for effectively exploring associations among DNA variation, transcriptomic profiles, and phenotypic diversity-especially in species with complex genomes^56,57^. We conducted RNA-seq analysis on shoot and root tissues from 110 wheat core accessions, uniformly distributed among three subgroups and representing abundant WUE_p_ variation within the population (**Supplementary Fig. 2a**), under WW, DS1 and DS2 conditions (**Fig. 1b and Supplementary Fig. 3a**).

We obtained 638 high-quality transcriptome datasets, yielding an average of 6.69 million uniquely mapped paired-end reads per sample (**Supplementary Table 3**). Principal component analysis (PCA) indicated that samples from WW, DS1, and DS2 conditions were clustered in a trend following the WW-DS1-DS2 in both root and shoot tissues, with a clear separation between tissues, suggesting the impact on gene expression is tissue > treatments > genotypes **(Fig. 2a and Supplementary Fig. 3a**). Gene expression levels generally remained stable, while expression coefficient of variation (CV) among accessions decreased under drought stress **(Fig. 2b and Supplementary Fig. 3b**). Significantly, expression fold-change levels and CV between drought-stress and well-water conditions (DS1/WW, DS2/WW) were higher than between drought gradients (DS2/DS1) **(Fig. 2c and Supplementary Fig. 3b**).

**Fig. 2.**
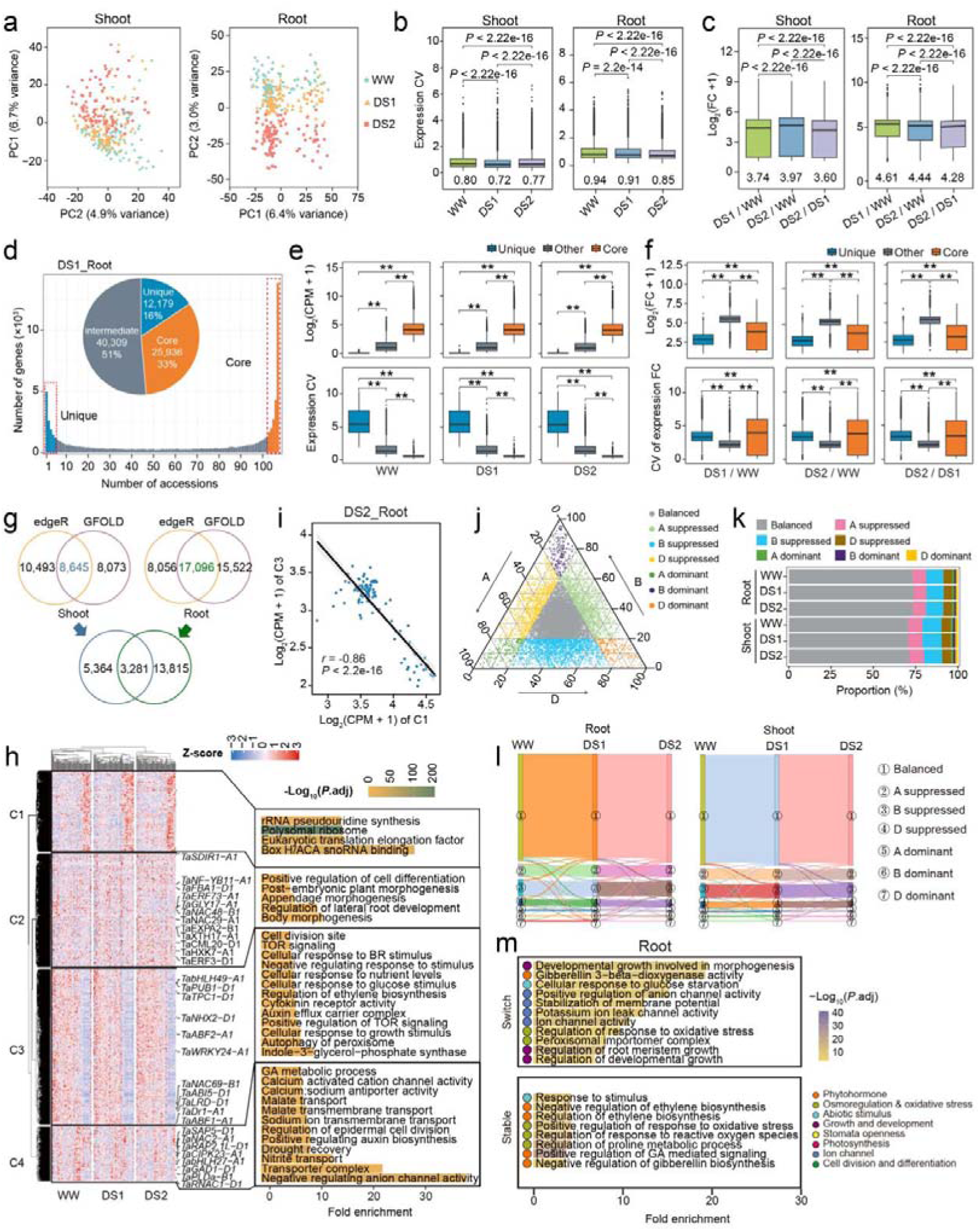
Pan-transcriptome dynamics of wheat responding to water scarcity. **(a)** Principal component analysis (PCA) of population-level transcriptome data of root (left) and shoot samples (right) under different conditions. Each dot represents a sample from each accession, with samples from each condition was indicated by different colors. **(b, c)** Comparison of the expression CV (e), and expression foldchange (f) among expressed genes in shoot (left) and root (right) under different conditions. In the box plot, borders represent the first and third quartiles, center line denotes median, and whiskers extend to 1.5 times the interquartile range beyond the quartiles. Wilcoxon rank sum test was used to determine the statistical significance between two groups. **(d)** The number of expressed genes detected in each accessions. The genes with CPM > 0.5 in at least one of the six samples were considered expressed genes. The genes expressed in > 90%, < 10%, and 10% ∼ 90% of the accessions were defined as core genes (in or orange), unique genes (in blue) and other genes (in gray), respectively. (**e, f**) Comparison of the expression level (upper panel in b), expression CV (lower panel in b), expression foldchange (upper panel in c), and expression foldchange CV (lower panel in c) among core, unique and other genes of root under different conditions. In the box plot, borders represent the first and third quartiles, center line denotes median, and whiskers extend to 1.5 times the interquartile range beyond the quartiles. Wilcoxon rank sum test was used to determine the statistical significance between two groups. **, *P* < 0.01. **(g)** Venn diagram displaying the overlap between population-wide differentially expressed genes (DEGs) identified in root and shoot samples. EdgeR and GFOLD were used to identify the population-wide DEGs. **(h)** Heatmap showing the k-means clustering of DEGs in root, with reported drought-resistance genes labeled, and the top enriched Gene ontology terms shown on the right. Fisher’s exact test were used for enrichment analysis. **(i)** The scatterplots showing the correlation of population-level average gene expression between C1 and C3 in root under DS2 condition. Each dot represents an accession. The black line represents the regression trend calculated by the general linear model, and the 95% confidence intervals were indicated with gray shade. The *r* value and *P* values were determined by Pearson correlation coefficient analysis. **(j)** Ternary plot showing homoeolog expression bias of triads. Only the triads with at least one DEGs and total expression levels with CPM > 0.5 were used. Each dot represents a gene triad with an A, B, and D coordinate consisting of the relative contribution of each homoeolog to the overall triad expression. The colors of the circle indicate different expression bias types. **(k)** Proportion of triads in each category of homoeolog expression bias across the tissue and water conditions. Only the triads with at least one DEGs and total expression levels with CPM > 0.5 were used. **(l)** Sankey diagram showing the state switching of triads in DEGs under different water conditions. Only the triads with at least one DEGs and total expression levels with CPM > 0.5 were used. **(m)** Enriched Gene Ontology terms for switch (upper panel) and stable (lower panel) triads in root. The length of shade indicates gene ratio enriched in each term, with color representing the - log_10_(*P*.adjust). Fisher’s exact test were used for enrichment analysis. Circles in the left indicate the functional group of GO terms.

Analysis of population-level transcriptomics revealed a comparable number of expressed genes among accessions, categorized into unique (expressed in <5% of accessions), core (>95%) and intermediate genes in root and shoot tissues based on expression frequency (CPM > 0.5) **(Fig. 2d**). Core genes exhibited the highest expression levels with minimal expression CV across accessions under varying water conditions, indicating high stability. In contrast, unique genes had lower expression level but significantly higher expression CVs **(Fig. 2e**), reflecting greater variability. Notably, core genes were more responsive to drought than unique genes, displaying considerable variation in expression CV among accessions, underscoring their role in phenotypic diversity in response to drought **(Fig. 2f**). Gene Ontology (GO) analysis further differentiated these gene groups: unique genes were enriched in ion channel activity, drought response, and developmental regulation, while core genes are primarily associated with enzymatic activity (**Supplementary Fig. 3c**). This suggests that core genes are likely essential for maintaining cellular homeostasis and exhibit high expression stability across wheat accessions. Conversely, the variability in unique genes underscore their essential role in accession-specific drought-induced phenotypic diversity and adaptive pathways.

A population-level transcriptomics analysis identified 8,645 and 17,096 differentially expressed genes (DEGs, see method for details) in shoot and root tissues, respectively, across treatment and accessions (**Supplementary Table 4**). Only 14.61% of DEGs were shared between tissues, implying distinct drought-response mechanisms in roots and shoots that contribute to phenotypic diversity **(Fig. 2g**). Most DEGs were identified under drought-stress versus well-watered conditions (DS1/WW, DS2/WW), with similar number of genes induced and repressed by drought (**Supplementary Fig. 3d**). Clustering of DEGs revealed four groups in both tissues, containing numerous known drought-resistance genes, such as *TaNHX2-D1*^58^ and *TaNAC48-B1*^13^ in root, and *TaERF87-A1*^59^ and TaSINA2B^36^ in shoot **(Fig. 2h, Supplementary Fig. 3e**). Notably, *TabHLH27-A1*^34^, which promotes both root and shoot biomass under well-limited conditions, was differentially expressed in both tissues. GO analysis revealed distinct enrichment patterns: C1 (translation), C2 (morphogenesis-related terms), C3 (phytohormone and stimuli response), and C4 (ion transport) **(Fig. 2h, Supplementary Fig. 3e, and Supplementary Table 5**). Interestingly, C1 expression negatively correlated with C3 across 110 accessions under all conditions, indicating an antagonistic relationship between growth and stress-response pathways **(Fig. 2i, Supplementary Fig. 3f**).

Subgenome-divergent regulation in allopolyploid wheat enhances genome plasticity and adaptability under stress^60–62^. We analyzed homoeologous expression bias across water conditions, categorising most triads (CPM >0.5) as balanced (72%–75%), followed by single-homoeolog suppressed (21%–24%), and single-homoeolog dominance (3%–4%) (**Supplementary Fig. 3g**). DEGs exhibited fewer balanced triads and more dominant and suppressed triads **(Fig. 2j, k**). With most triads remained stable, a small fraction shifted expression pattern across conditions (7% in shoots, and 9% in roots) (**Fig. 2l and Supplementary Table 6**). Stable triads were associated with phytohormone, osmoregulation and oxidative stress (**Fig. 2m and Supplementary Fig. 3h**). In contrast, switch triads displayed tissue specific enrichment: roots showed enrichment in ion channel activity, phytohormone signaling, and growth regulation, while shoots highlighted photosynthesis and phytohormone related terms **(Fig. 2m and Supplementary Fig. 3h**). These findings suggest that wheat adapts to drought by modulating sub-genome expression bias, enabling tissue-specific response to environmental stress.

Our pan-transcriptome analysis reveals distinct drought responses in wheat, with core genes maintaining homeostasis and unique genes driving diversity. Differentially expressed genes show tissue-specific adaptations, while subgenome-divergent regulation highlights balanced and adaptive shifts in triads.

### Mapping and characterizing eQTL hotspots driving wheat’s drought resilience

Phenotypic diversity is usually determined by the variability in expression of related genes, and thus it is important to explore causal genetic variations in modulating gene differential expressions^63,64^. To pinpoint the regulatory elements underlying complex traits, we conducted an expression quantitative trait locus (eQTL) mapping using SNPs in the 110 core accessions (from Wheat660K SNP array and Whole-exome sequencing, see method for details), uncovering 146,966 eQTLs associated with 8,218 Genes (eGene) across the wheat genome (**Supplementary Fig. 4 and Supplementary Table 7**). These eQTLs averaged a spacing of 0.2 Mb and were mainly distributed across distal intergenic regions, exons, and promoter regions within 1 kb of transcription start site **(Fig. 3a, b and Supplementary Fig. 5a**). By clustering tightly linked eQTLs (with average size of 1.91 kb), we identified distinct regulatory loci, or pan-regulons, separated into *cis*-eQTLs and *trans*-eQTLs based on LD distance threshold of 2.4-Mb from their respective eGenes **(Fig. 3c and Supplementary Fig. 5b**). Notably, 27%–28% of eGenes were controlled by *cis*-eQTLs, 23%–25% by *trans*-eQTLs, and nearly half by a combination of both **(Fig. 3d**). Consistent with prior research^45,65^, *cis*-eQTL exhibited a larger impact on gene expression variability compared to *trans*-eQTL **(Fig. 3e**). The *trans*-eQTLs were further categorized into intra-chromosomal (eQTL and eGenes located in the same chromosome) and inter-chromosomal (eQTL and eGenes located in different chromosomes), with B-D subgenomic interactions being predominant, while A-A interaction was least frequent **(Fig. 3c and Supplementary Fig. 5b**), suggesting a subgenomic bias in expression regulation.

**Fig. 3.**
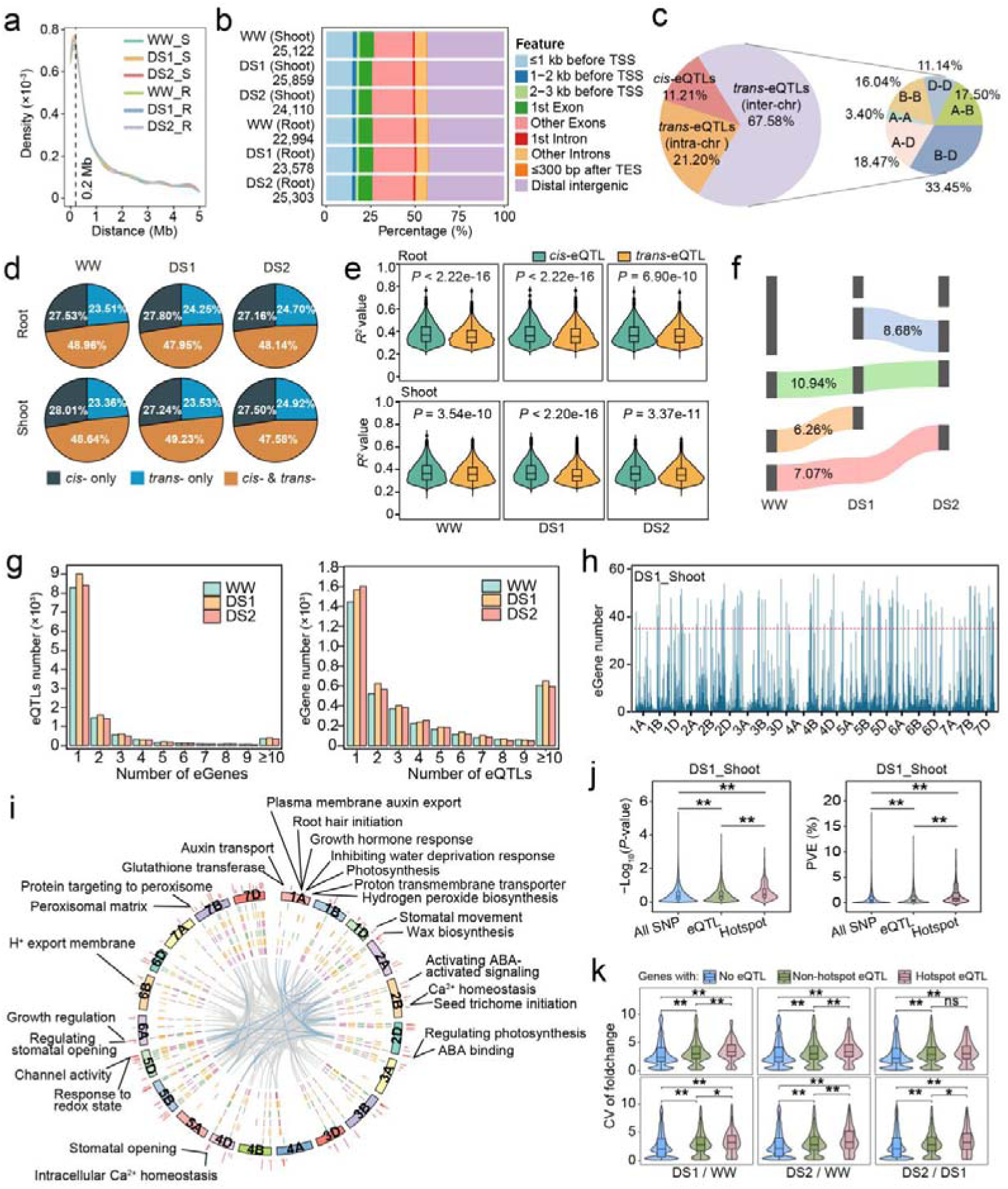
Pan-regulons modulating gene expression diversity under water scarcity. **(a)** Density distribution of distances between eQTLs and their regulated eGenes. The dashed vertical line indicates the average eQTL-eGene distances by pooling regulatory relationships of each tissue and water condition. **(b)** Distribution of eQTLs relative to genes. **(c)** Proportion distribution of *cis*- and *trans*-eQTLs and the subgenomic interactions involved in *trans*-eQTL regulation in root tissue. The inter-eQTLs indicates that eQTL and eGenes located in the same chromosome, and intra-eQTLs means that the eQTL and its eGenes are on different chromosomes. **(d)** Category proportion of eGenes according to the eQTLs type. **(e)** Comparison of the *R^2^* value of eQTL between *cis*- and *trans*-eQTLs under different conditions. In the box plot, borders represent the first and third quartiles, center line denotes median, and whiskers extend to 1.5 times the interquartile range beyond the quartiles. Wilcoxon rank sum test was used to determine the statistical significance between two groups. **(f)** Sankey diagram showing the sharing and specificity of eQTLs in root among water conditions. The numbers indicated proportion of eQTLs of each group relative to the total eQTL number. **(g)** Distribution of eQTLs with each number of target genes (left), and eGenes with each number of eQTLs (right) in root across different water conditions. **(h)** Number of eGenes regulated by distal eQTLs in DS2_Root condition. The horizontal dashed line indicates the threshold of eGene numbers regulated by eQTL hotspot regions. **(i)** The genome identification of eQTL hotspots under each tissue and condition. The most outside circle, chromosomes names; Sequentially from outer to inner layers: eQTLs from WW_Shoot, DS1_Shoot, DS2_Shoot, WW_Root, DS1_Root and DS2_Root. The most inside layer, distal regulatory relationships between eQTLs and eGenes. Drought-resistant-related GO terms of some eQTL hotspots were indicated around the circle graph. **(j)** Comparison of -log_10_(*P*-value) and phenotypic variation of SNPs explained in the GWAS of WUE_p_ among all GWAS SNPs, non-hotspot eQTLs and hotspot eQTLs. The non-hotspot eQTLs and hotspot eQTLs in DS2_Root were used. Wilcoxon rank sum test was used to determine significant differences. *, *P* < 0.05; **, *P* < 0.01. **(k)** Comparison of the CV of expression foldchange among all GWAS SNPs, non-hotspot eQTLs and hotspot eQTLs. Wilcoxon rank sum test was used to determine significant differences. ns, no significance; *, *P* < 0.05; **, *P* < 0.01.

To identify regulons that govern drought-specific gene expression changes, we analyzed eQTL stability across WW and DS1, DS2 conditions. We found that most eQTLs were condition-specific, with only ∼11% remaining static across all conditions **(Fig. 3f and Supplementary Fig. 5c**), highlighting the regulatory specificity for drought responses. Dynamic eQTLs were more prevalent in the A-subgenome, while the D-subgenome contributed more to static eQTLs (**Supplementary Fig. 5d**), aligning with previous findings on subgenome specialization in wheat^60,62^. Most eGenes were regulated by a single eQTL, yet we identified numerous eQTL hotspots with multiple eGene targets (see method for details) **(Fig. 3g, h and Supplementary Fig. 5e, f**). The eGenes of hotspots preferred to locate on the D-subgenome (**Supplementary Fig. 5g**), align with the critical role of D-subgenome in conferring abiotic stress resistance^66,67^. A sizable proportion of these hotspots overlapped with known drought-resistance genes and GWAS signals, and their associated eGenes were enriched in pathways critical for drought adaptation, including regulation of water loss (e.g., “stomatal opening” and “wax biosynthesis”), water deprivation response (e.g. “inhibiting water deprivation”), osmoregulation, oxidative stress (e.g., “hydrogen peroxide biosynthesis”, “protein targeting to perxisome”), phytohormone (for example, “auxin transport” and “activating ABA-activated signaling”), and photosynthesis **(Fig. 3i**). eQTL hotspots had higher detection power and phenotypic impact in GWAS analysis than non-hotspot SNPs, with non-eQTL SNPs showing the lowest detection power and effect sizes **(Fig. 3j and Supplementary Fig. 5h**). Moreover, eGenes regulated by hotspots displayed significantly higher expression level and variability across accessions in response to drought, underscoring their role in driving gene expression diversity under stress **(Fig. 3k and Supplementary Fig. 5i**).

Collectively, we conducted eQTL mapping in 110 wheat accessions, identifying 146,966 eQTLs linked to 8,218 genes. Dynamic eQTLs, particularly in the A-subgenome, were drought-specific. eQTL hotspots enriched in drought-related pathways showed stronger GWAS signals and phenotypic impacts.

### Identification of gene modules and key regulators in wheat WUE and drought resistance

To investigate how coordinated gene expression contributes to phenotypic variation in WUE and drought resistance, we performed a weighted gene co-expression network analysis (WGCNA) on pan-transcriptome DEGs. This analysis clustered genes into 22 modules in shoots and 20 modules in roots after merging highly similar modules, each marked by a unique color **(Fig. 4a. Supplementary Fig.6a and Supplementary Table 8**). Most modules contained fewer than 1,000 genes (**Supplementary Fig. 6b**), and each displayed distinct patterns of relative expression under varying water conditions (**Supplementary Fig. 6c, d**), indicating possible roles in drought response. Correlations between module and four key traits identified 15 modules in shoot and 13 modules in root significantly associated with at least one trait (**Supplementary Fig. 6b**). For further analysis, we focused on the top three modules for each trait, yielding five modules (Blue, Lightgreen, Grey60, Lightyellow, and Red) in roots. GO enrichment analysis revealed that these modules were enriched for GO terms related to water stress response, phytohormone signaling, root development, osmoregulation, and oxidative stress metabolism **(Fig. 4b**). Similarly, top modules identified in shoots (Darkred, Tan, Yellow, Royalblue, Grey60, Salmon, Greenyellow, and Midnightblue) were enriched for photosynthesis, phytohormone regulation, osmotic stress response, and oxidative stress terms (**Supplementary Fig. 6e, f**).

**Fig. 4.**
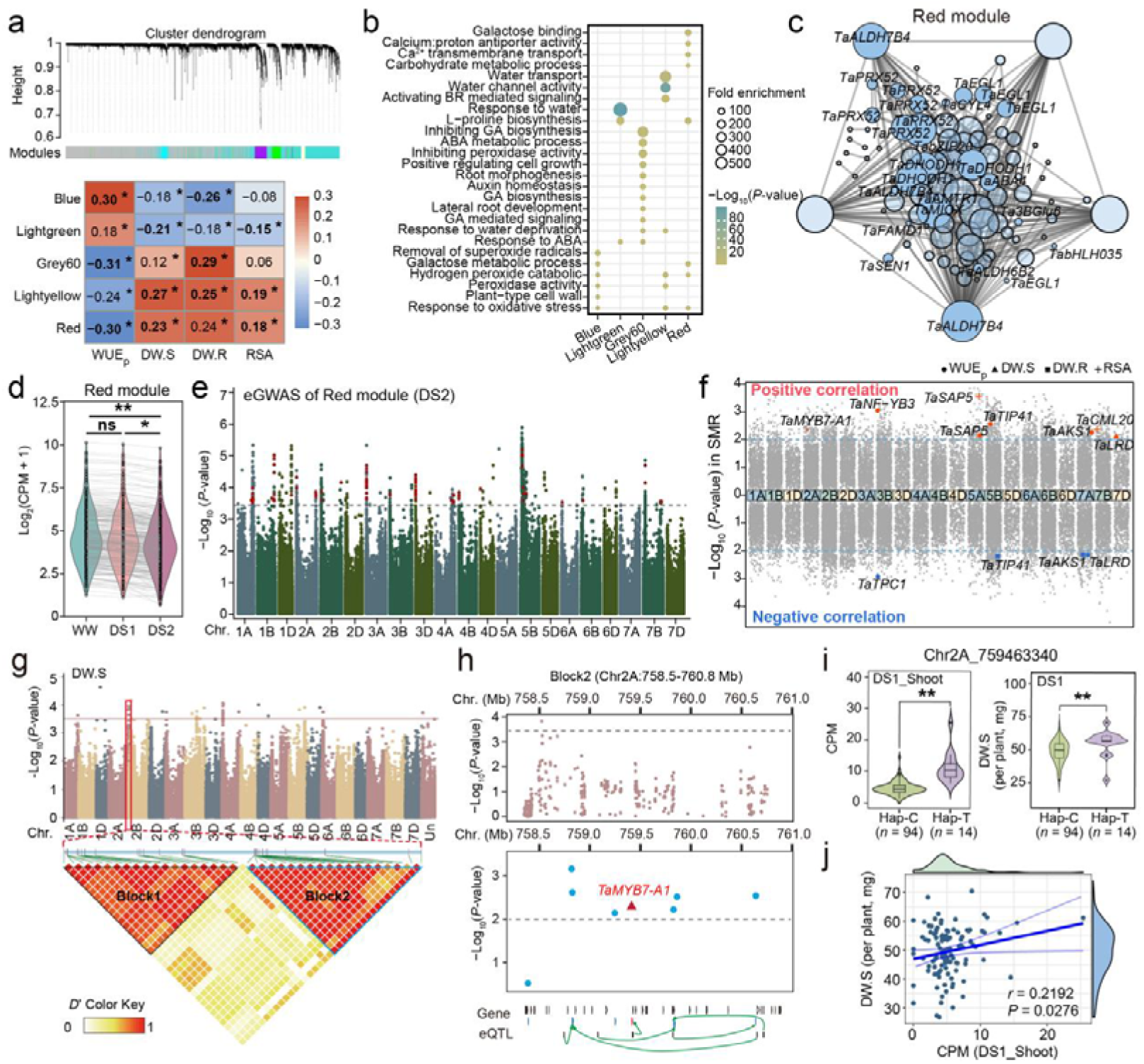
WGCNA and SMR prioritize regulators for WUE and drought-resistance in wheat. **(a)** Hierarchical cluster dendrogram showing co-expressed modules (upper panel) in root and top correlated modules with each trait (lower panel). The co-expressed modules were identified by weighted gene co-expression network analysis and each module was indicated with a branch, and each gene was indicated with a leaf with ordinate showing the clustering distance of each gene. Top three correlated modules of each trait were shown in the heatmap, with different colors showing the Pearson correlation coefficient (values displayed in the square, asterisks indicated the significant correlation with *P* < 0.05). **(b)** Enriched Gene Ontology (GO) terms for top correlated modules in (a). The size of dots indicates gene ratio enriched in each term, with color representing the *P*.adjust value. Fisher’s exact test were used for enrichment analysis. **(c)** Co-expression gene network in the Red module. The size of the cycles indicates number of linked genes in the module. **(d)** Comparison of the expression level of Red module in root among three water conditions. Each gene in the module is plotted as a dot, and their expression levels under different water conditions are linked with a light gray line. Wilcoxon rank sum test was used to determine significant differences. ns, no significance; *, *P* < 0.05; **, *P* < 0.01. **(e)** Manhattan plots illustrating the eGWAS Llog_10_(*P*) values using the expression level of the Red module as a trait. Horizontal dashed line indicates the genome-wide significant threshold of Llog_10_(*P*-value) = 3.50. The SNPs identified as the eQTL in root were marked in red. **(f)** Manhattan plot showing the integrative results of SMR for four traits. The SMR scores (y-axis) are plotted against gene positions (x-axis) on each of the chromosomes. The dashed lines indicate the threshold of -log_10_(*P*-value) = 2.0 for defining SMR significant genes in the following analysis. Known genes related to drought-resistance or WUE are labeled, and marked as red dots if their expression level positive correlated with the phenotypic data, or blue dots if negative correlated. **(g)** Manhattan plot illustrating the Llog_10_(*P*-value) values in the GWAS of DW.S, with a heatmap displaying linkage disequilibrium (LD) between SNPs within the 4.8-Mb physical interval flanking the peak SNPs on chromosome 2A. In the Manhattan plot, y-axis indicates Llog_10_(*P*-value) values of each SNP, x-axis indicates SNP position on each chromosome, and the horizontal dashed line indicates the genome-wide suggestive statistical-significance threshold of -log_10_(*P*-value) = 3.50. The LD heatmap intensity of color from white to black represents the range of *r^2^* values from 0–1. **(h)** Local Manhattan plot showing significant SNPs in GWAS (upper panel), SMR-correlated genes (middle panel), and eQTL-eGene regulatory relationships (lower panel) in the LD block2 in (g). Y-axis, the Llog_10_(*P*-value) values in the GWAS (in upper panel) or SMR (in middle panel) of DW.S. X-axis, the chromosomes location. Horizontal dashed lines indicate the genome-wide suggestive statistical-significance threshold (upper panel, -log_10_(*P*-value) = 3.50 in GWAS; middle panel, -log_10_(*P*-value) = 2.00 in SMR). The *TaMYB7-A1* is denoted in red. **(i)** Comparison of the expression level (left) and DW.S (right) of different haplotypes defined by the SNP chr2A_759463340 among wheat accessions. SNP chr2A_759463340 was the eQTL regulates *TaMYB7-A1* expression. Wilcoxon rank sum test was used to determine significant differences. **, *P* < 0.01. **(j)** Scatterplot showing the correlation of *TaMYB7-A1* expression and DW.S under DS1 condition. Each dot in the main plot represents data of an accession, and marginal distribution plots of the CPM and DW.S were plotted above and right of the main plot, respectively. The blue line represents the regression trend calculated by the general linear model, and the 95% confidence intervals were indicated with lightblue curves. The *r* value and *P* values were determined by Pearson correlation coefficient analysis.

Among these, the Lightgreen, Lightyellow, and Red modules of root showed significant correlation with all four traits **(Fig. 4a**). Notably, the Red module (size = 314) was negatively correlated with WUE_p_ (*R* = -0.30) but positively associated with other three traits (*R* = 0.18–0.24). This module was enriched in genes involved in osmolyte accumulation and oxidative stress, suggesting a role in regulating WUE_p_ by enhancing osmotic and hypoxic stress tolerance **(Fig. 4a, b**). A network map of interactions within the Red module revealed hub genes with known roles in osmoregulation, such as *TaALDH7B4*, *TaDHOD1*, *TaPRX52* and key transcription factors *TaEGL1*, *TabZIP20*, *TaABA4*, and *TabHLH035* **(Fig. 4c**). Expression of Red module genes was notably reduced under water-deficit conditions, especially in DS2 **(Fig. 4d**). To identify genomic loci distally regulating this module, we conducted a GWAS using the expression pattern of Red module as a trait (eGWAS), revealing 77 genomic loci, with 29 loci (37.66%) overlapping with eQTLs under DS2 **(Fig. 4e**). Motif analysis of DNA regions flanking these loci showed enrichment of binding sites of ERF, C_2_H_2_, and LBD transcription factors, indicating potential regulatory control by these TFs (**Supplementary Table 9**). Similarly, network analysis of the Darkred module in shoots highlighted aquaporin TaNIP5;1 and dehydration-responsive RD22, potentially regulated by ERF, NAC, and C_2_H_2_ TFs (**Supplementary Fig. 6g-i and Supplementary Table 9**).

To prioritize candidate causal genes for drought tolerance and WUE, we applied a Summary data-based Mendelian Randomization (SMR) test. This test identified 830, 473, 464, and 324 genes significantly associated (*P* < 0.01) with WUE_p_, DW.S, DW.R, and RSA (**Supplementary Table 10**), respectively. Among these, nine genes with known role in drought resistance were positively associated with one or more traits, while four were negatively associated (**Fig. 4f**). One notably gene, *TaMYB7-A1* (homolog of *OsMYB6*), emerged as a strong candidate due to its known role as a positive regulator in drought and salt stress tolerance in rice^68^. This gene resides within a 2.3-Mb LD block (Block2, chr2A: 758.5–760.8 MB) associated with DW.S **(Fig. 4g, h**). Of the 44 genes within this region, only *TaMYB7-A1* was both differentially expressed in the pan-transcriptome and identified as an SMR-associated gene for WUE-related traits **(Fig. 4h**). To further investigate, we analyzed regulatory variants of *TaMYB7-A1* and identified a SNP downstream of the gene acting as a proximal regulatory eQTL. Accessions carrying the Hap-T variant (*n* = 14) exhibited higher *TaMYB7-A1* expression and DW.S than those with the Hap-C variant (*n* = 94) **(Fig. 4i**). Given that *TaMYB7-A1* expression positively correlates with DW.S **(Fig. 4j**), this gene is a strong candidate for contributing to phenotypic variation in WUE across the population.

We conducted a WGCNA on wheat pan-transcriptome DEGs, identifying key modules linked to drought response and WUE. Significant modules such as Red were correlated with multiple traits, including WUE, osmotic stress tolerance, and oxidative stress. A GWAS revealed loci regulating these modules, with 29 loci overlapping with eQTLs under drought stress. SMR analysis identified 830 genes associated with WUE-related traits, among which TaMYB7-A1 emerged as a key candidate, positively correlating with drought tolerance and shoot dry weight (DW.S).

### Integrative omics analysis elucidates the genetic basis underlying wheat WUE

To identify key regulators and causal genetic variants for WUE in wheat, we constructed drought-resistance and WUE associated network based on the co-localization of LD regions surrounding GWAS signals (GWAS bins) with SMR genes. This model provides a functional network of drought adaptation regulated by genetic variations, encompassing 146 GWAS bin, 333 SMR genes, and 338 eQTL regions modulating their expression, with 7 eQTLs and 35 genes overlapping across networks of multiple traits **(Fig. 5a, Supplementary Fig. 7a and Supplementary Table 11**). Among the SMR genes within the WUE_p_ network, several are known to play roles in water deprivation, osmotic stress, water transport, ABA response, stomatal movement, wax biosynthesis, calcium signaling, and growth processes in rice or *Arabidopsis thaliana* **(Fig. 5a, b**). Of these, 11 genes are regulated by proximal eQTLs (within their GWAS bins), while 19 are modulated by distal eQTLs (outside their GWAS bins).

**Fig. 5.**
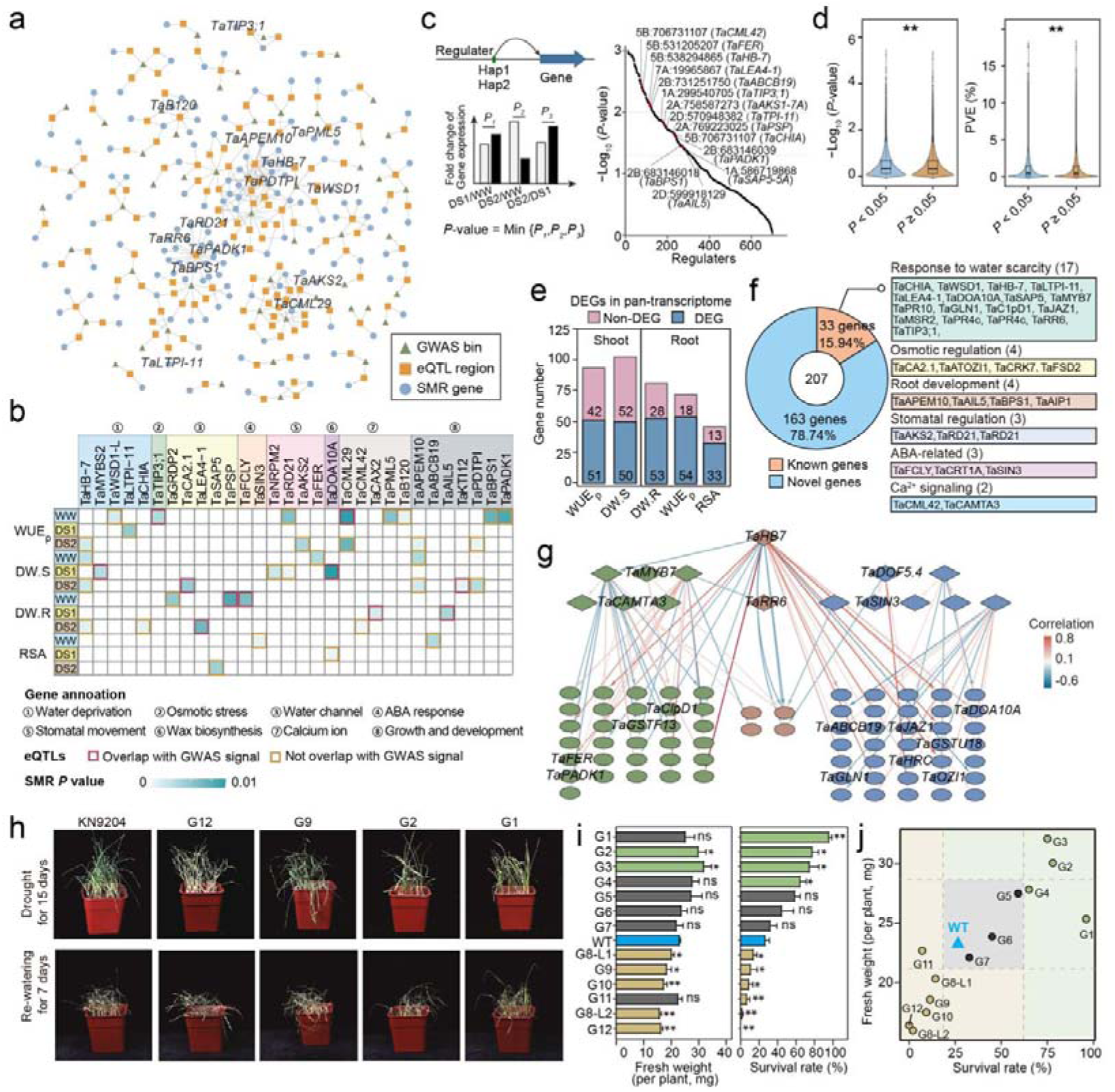
Integrative omics prioritize candidate causal genes for drought-resistance in wheat. **(a)** A genetic network of WUE_p_ was constructed by integrating GWAS signal, eQTL and SMR genes. The GWAS, eQTL and SMR data of WUE_p_ under different conditions were pooled. An SMR gene located within the LD region of a GWAS signal, or an SMR gene with an eQTL located within this LD region, was considered co-localized and linked to the GWAS signal. The GWAS signal, eQTL and SMR genes were indicated with different symbols, and the line between SMR gene and eQTL indicates a regulatory relationship, and lines between GWAS signals and SMR genes or eQTLs indicate co-localization of LD regions. **(b)** The overlapping of GWAS signal and eQTL with 30 SMR genes whose homoeolog gene in rice or *Arabidopsis thaliana* were reported to involved in drought-resistance or WUE. The 30 genes were divided into eight functional categories (marked in different colored shades). The SMR genes co-localized with GWAS signals of each trait were marked by square with color according to its SMR *P*-value with this trait. The SMR gene whose eQTL co-localized with GWAS signals of each trait were marked in red frame, while these not co-localized with GWAS signals were in brown. **(c)** A ranking scatter plot showing the pan-regulons significantly alters target genes’ response to drought. As illustrated in the schematic (left panel), an eQTL divide the population into two groups carrying different haplotype. The significance in the differences of these two groups’ response to drought stress were calculated, and the minimum value among *P*-values for DS1/WW, DS2/WW, and DS2/DS1 were retained. The eQTLs above the horizontal dashed line (statistical-significance threshold of *P*-value = 0.05) were considered as pan-regulons. The pan-regulons identified as eQTLs for drought-resistance or WUE related genes were in red dots and labeled. **(d)** Comparison of -log_10_(*P-*value) and phenotypic variation of SNPs explained in GWAS between the eQTLs above and below the horizontal dashed line (*P* < 0.05 and *P* ≥ 0.05, respectively) in (c). For each eQTL, the highest -log_10_(*P*-value) value in the GWAS of multiple traits were used. Wilcoxon rank sum test was used to determine significant differences. **, *P* < 0.01. **(e)** Bar graphs showing the gene number identified in the genetic network of each trait. Among which, genes identified as a DEG in the population-level transcriptome data were considered as candidate genes and marked in blue. **(f)** Proportion of the candidate genes in (e) of each category. The symbol of known genes are listed according to its functional category. **(g)** A hierarchical regulatory network of 207 candidated genes. Diamonds, TFs; Ovals, functional genes. Red and blue arrows indicate the positive and negative regulation relationship, respectively, with darker the color, the stronger its correlation. The shoot-specific, root-specific, and shared genes were in green, blue, and brown, respectively. **(h)** Representative photographs showing the phenotype of wild-type KN9204 (WT) and several loss-of-function mutant lines under drought stress. Photos (each biological replicate with 36 seedlings) were taken after water-holding for 15-days (upper panel) and then water recovery for 7-days (lower panel). **(i)** Bar graphs showing the fresh weight (lift) and survival rate (right) of WT and loss-of-function mutant lines of candidate genes. The fresh weight and survival rate were investigated after water recovery for 7-days. Data are the mean ± S.D. of four biological replicates. Student’s *t* test were used to determine the statistical significance between each mutant line and the WT. ns, no significance; *, *P* < 0.05; **, *P* < 0.01. Lines with higher fresh weight or survival rate compared to WT were marked in green, whereas these with lower fresh weight or survival rate were in brown. The lines without statistical significance with WT were marked in gray, and the WT was highlight in blue. **(j)** Scatter plot showing the survival rate and fresh weight of WT and each mutant line. Two horizontal and vertical dashed lines were used to divide accessions with/without significant differences in fresh weight and survival rate, respectively, compared to WT. By combining the phenotypes of two traits, lines with higher and lower drought-resistance compared to WT were marked in green and brown, respectively. The lines without statistical significance with WT were marked in gray, and the WT was highlight in blue.

To further reveal the genetic variations that diverse drought responses among wheat accession, we examined the effects of regulatory sites on gene expression under different conditions. DNA variation at 318 regulators influenced the expression response of downstream genes to drought stress, as illustrated by a distal regulatory site at chr5B:706731107, which modulates *TaCML42* expression **(Fig. 5c**). Importantly, these regulatory variants exhibited higher detection power in GWAS **(Fig. 5d**), underscoring their relevance to WUE_p_ traits. We refined candidate genes for drought response by intersecting SMR genes with DEGs identified in the pan-transcriptome, yielding 207 candidates **(Fig. 5e, f**). Among these, 33 genes (15.94%) are wheat homologous of genes involved in water scarcity response (e.g., *TaMYB7-A1*^68^), osmotic regulation (e.g., *TaCA2.1*^69^), root development (e.g., *TaBPS1*^70^), stomatal regulation (*TaAKS2*^71^), and ABA-related genes (e.g., *TaSIN3*^72^). A hierarchical regulatory network of these 207 genes included six TFs regulating gene expression in shoot, seven TFs regulating gene expression in root, and two TFs active in both shoot and root, reflecting a complex inter-regulatory module controlling WUE-related traits in wheat **(Fig. 5g**).

To validate the functional relevance of these candidate genes in drought resistance, we assessed fresh weight (FW) and survival rate (SR) in EMS-mutagenized homozygous mutant lines of the wheat varieties KN9204^50^. We analyzed 13 mutant lines of 12 random selected candidate genes (with two lines for gene G8) alongside wild-type KN9204 in drought conditions **(Fig. 5h, Supplementary Fig. 7b and Supplementary Table 12**). Mutants of two genes exhibited increased FW, while five lines for four genes showed reduced FW compared to the wild-type KN9204; mutants for four genes had higher SR, while mutants for five genes (six lines) had lower SR **(Fig. 5i**). In total, mutation in four genes enhanced drought-resistance (increased FW or SR), while mutation in five genes diminished drought-resistance **(Fig. 5j**). This indicates that the majority (9 out of 12 genes, 75%) of the tested candidates significantly influence FW or SR under drought, confirming the validity of our strategy for identifying drought-resistance regulators in wheat.

In conclusion, our multi-omics approach, integrating GWAS, eQTL, SMR, and pan-transcriptome data, identified 207 candidate causal genes as key regulators for WUE-related traits in wheat. Functional assessment of candidates in gene-indexed mutants demonstrated their significant roles in modulating drought resistance.

### Overexpression of *TaMYB7-A1* enhances drought-resistance and WUE in wheat

To investigate the causal role of systemic identified factors, we selected *TaMYB7-A1* (*TraesCS2A02G554200*) for functional analysis. Previously, *TaMYB7-A1* was over expressed in wheat^73^, and here we examine its effects on drought resistance. We analyzed DNA variations within ∼10-Kb genomic region flanking *TaMYB7-A1* across the wheat population and identified 27 SNPs. The population was divided into three haplotype groups: Hap-1 (13 accessions), Hap-2.1 (46 accessions), and Hap-2.2 (47 accessions) **(Fig. 6a)**. Hap-1 accessions showed significantly higher *TaMYB7-A1* expression level and greater drought tolerance (DW.S) compared to Hap-2 and Hap-3, which showed no significant differences **(Fig. 6b**). Thus, Hap-1 was considered the superior haplotype for drought resistance. Under water-limited (WL) conditions, *TaMYB7-A1* expression was significantly up-regulated in both shoot and root in Fielder (Hap-1), supporting its role in drought tolerance **(Fig. 6c**).

**Fig. 6.**
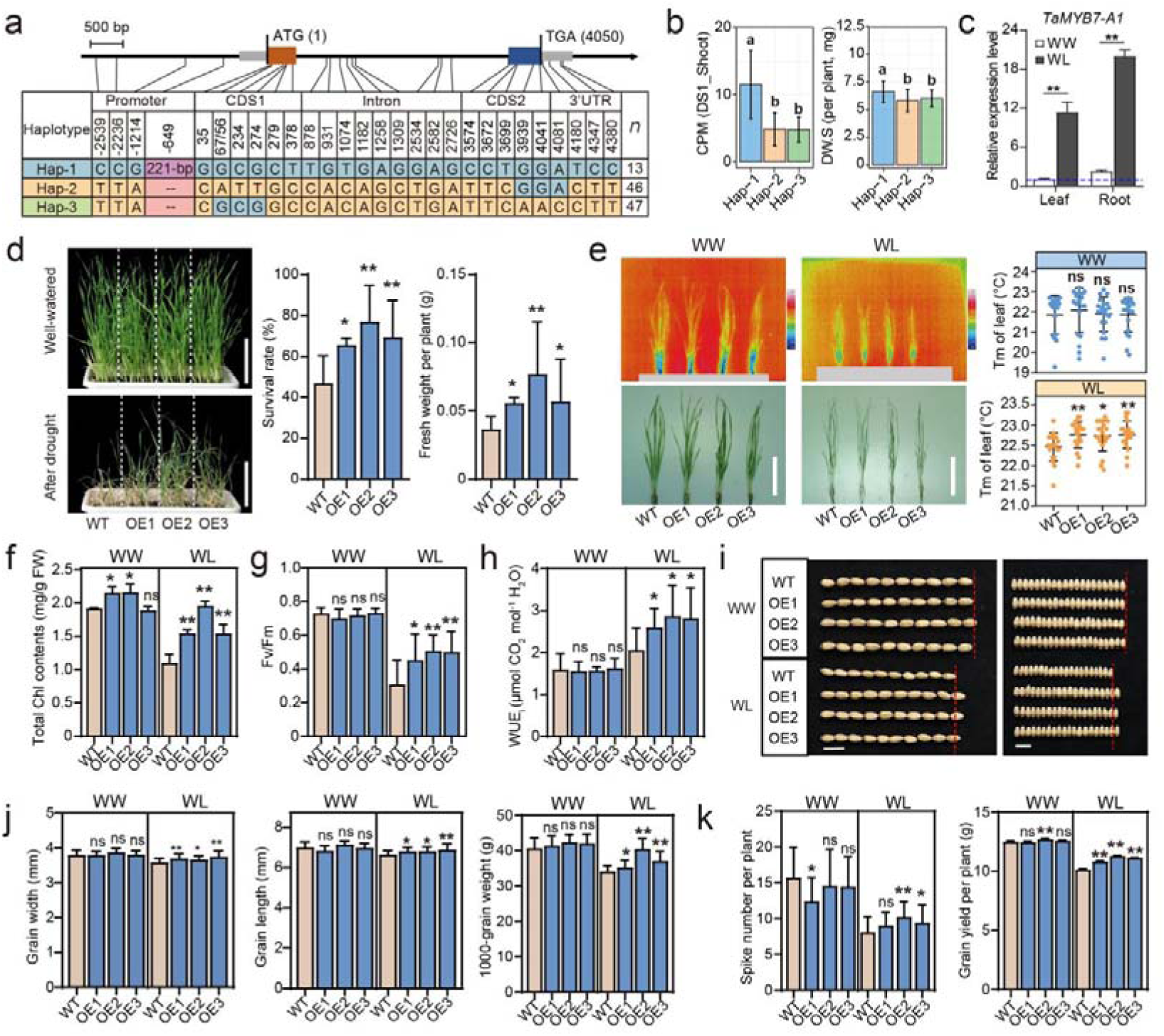
TaMYB7-A1 improves drought-resistance and WUE in wheat. **(a)** Schematic diagram showing the polymorphisms for each haplotype of *TaMYB7-A1* in the accessions used for pan-transcriptome. SNPs were identified by Affymetrix Wheat660K SNP arrays and Whole Exome Sequencing, and a 221-bp insertion/deletion was genotyped using specific primers described as Wang et al., 2024. **(b)** Comparison of the *TaMYB7-A1* expression level and DW.S among three haplotypes. DW.S and *TaMYB7-A1* expression level (DS1_shoot) in the pan-transcriptome accessions were used for analysis. Tukey’s HSD multiple comparison test was used to compare the Llog_10_(*P*-value) value between each group, with different letters indicate a statistically significant difference at *P* < 0.05. **(c)** The *TaMYB7-A1* expression pattern in shoot and root of seedlings under WW and WL conditions. RT-qPCR was used to determine the *TaMYB7-A1* expression pattern, with *TaTublin* as the internal control. The expression levels were normalized with WT under WW condition set to 1.0. Data are means ± S.D. of three independent biological replicates. Student’s *t* test were used to determine the statistical significance. **, *P* < 0.01. **(d)** Assessment of the survival rate and fresh weight per plant of Fielder and OE lines under severe drought stress. Representative photographs of three independent biological replicate (each with 28 seedlings; Scale bars, 10 cm) were taken and phenotypic data investigated after a 7-d period of recovery post-drought treatment. Data are means ± S.D. of three independent biological replicates. Student’s *t* test were used to determine the statistical significance between each OE line and Fielder. *, *P* < 0.05; **, *P* < 0.01. **(e)** Leaf temperature of Fielder and OE lines under WW and WL conditions investigated using infrared thermography. In the infrared imaging photographs (upper panel), red to blue colors indicate temperatures from high to low. The leaf surface temperature were quantitatively measured with randomly selected 20 image points from seedlings of each genotype. Data are means ± S.D. (*n* = 20). Student’s *t* test were used to determine the statistical significance between each OE line and Fielder. ns, no significance; *, *P* < 0.05; **, *P* < 0.01. Scale bars, 5 cm. (**f-h**) Comparison of the total Chl contents (f), Fv/Fm values (g) and WUE_i_ (h) between Fielder and OE lines under WW and WL conditions. Data are means ± S.D. (for total Chl contents, *n* = 4 independent biological replicates; for Fv/Fm and WUE_i_, *n* = 8 independent biological replicates). Student’s *t* test were used to determine the statistical significance between each OE line and Fielder. ns, no significance; *, *P* < 0.05; **, *P* < 0.01. **(i)** Representative photographs showing the grain length and width of Fielder and OE lines under WW and WL conditions in field. Red vertical lines indicates the length or width of ten Fielder grains. Scale bars, 1 cm. (**j, k**) Comparison of the grain legth, grain width, 1000-grain weight, spike number per plant and grain yield per plant between Fielder and OE lines under WW and WL conditions in field. Data are means ± S.D. (for grain width, grain length, 1000-grain weight and and grain yield per plant, *n* = 6 independent biological replicates; for spike number per plant, *n* = 40 independent biological replicates). Student’s *t* test were used to determine the statistical significance between each OE line and Fielder. ns, no significance; *, *P* < 0.05; **, *P* < 0.01.

The generated independent overexpression (OE) lines of *TaMYB7-A1^Hap-1^*were driven by the Ubiquitin promoter in wheat *cv.* Fielder^73^. Three lines with exhibited comparable over-expression levels in roots and shoots (**Supplementary Fig. 8a**), were evaluated under both well-watered (WW) and water (WL) conditions. Under WW conditions, no phenotypic differences were observed between OE lines and wild-type Fielder (WT). However, under WL condition, OE lines exhibited higher survival rates and fresh weight per plant to WT controls **(Fig. 6d)**. Additionally, OE lines developed larger root systems, with significantly increased root length, volume and surface area, under both WW and WL conditions (**Supplementary Fig. 8b**).

To assess leaf function, transpiration and photosynthesis were measured. While leaf surface temperatures were similar under WW conditions, OE lines exhibited higher leaf temperature under WL conditions, suggesting that *TaMYB7-A1* overexpression reduced water loss through transpiration **(Fig. 6e**). OE lines also retained higher chlorophyll levels especially under WL conditions, despite drought-induced declines **(Fig. 6f**). Furthermore, OE lines maintained significantly higher photosystem II functionality (Fv/Fm value from chlorophyll fluorescence measurements) than WT control plants under WL condition, but no change under WW condition (**Fig. 6g**). Collectively, reduced transpiration and increased photosynthesis improved WUE_i_ (the ratio of instantaneous net CO_2_ absorption rate to the transpiration rate) in OE lines compared to WT control plants under WL condition (**Fig. 6h**).

To evaluate the effects of TaMYB7-A1 on yield-related traits, we assessed OE lines under WW and WL field conditions. Under WW conditions, the OE lines exhibited no significant phenotypic differences from WT control plants in grain length, width, and weight **(Fig. 6i, j**). While OE2 displayed slightly fewer spikes per plant and OE3 demonstrated an increased spikelet number per spike, these variations did not affect overall grain yield, though OE2 showed a modest yield increase (**Fig. 6k and Supplementary Fig. 8c**). Under WL conditions, all three OE lines produced more spikes per plant, more spikelets per spike, and larger grains, leading to significantly higher grain yields **(Fig. 6k and Supplementary Fig. 8c**).

These findings indicate that *TaMYB7-A1* overexpression enhances drought resistance, WUE, and yield under water-limited conditions without compromising growth or yield under well-watered conditions, highlighting its potential for improving wheat performance in drought-prone environments.

### TaMYB7-A1 enhances wheat drought tolerance and WUE by regulating antioxidants, osmotic balance, and root development

To explore the molecular mechanisms by which TaMYB7-A1 regulates drought resistance and WUE in wheat, we conducted RNA sequencing on WT and *TaMYB7-A1* OE lines under WW and WL conditions (**Supplementary Fig. 9**). PCA revealed a clear separation of samples from WW and WL, indicating effective drought induction and significant impacts on gene expression **(Fig. 7a**). RNA-seq revealed 30,513 and 15,305 drought response genes (DRGs, WL compared to WW, fold changes ≥ 2 and adjusted *p*-value < 0.05) in root and shoot of WT control plants, respectively **(Fig. 7b and Supplementary Fig. 10a**). Drought-induced genes (up-regulated in WL conditions) in both root and shoot were enriched for GO terms related to “Response to water deprivation”, “Response to abiotic stimulus”, and “Response to hydrogen peroxide”. However, drought-suppressed genes (down-regulated in WL conditions) showed distinct patterns: shoot DRGs were enriched for photorespiration and senescence, while root DRGs were linked to oxidative stress (**Supplementary Fig. 10b**).

**Fig. 7.**
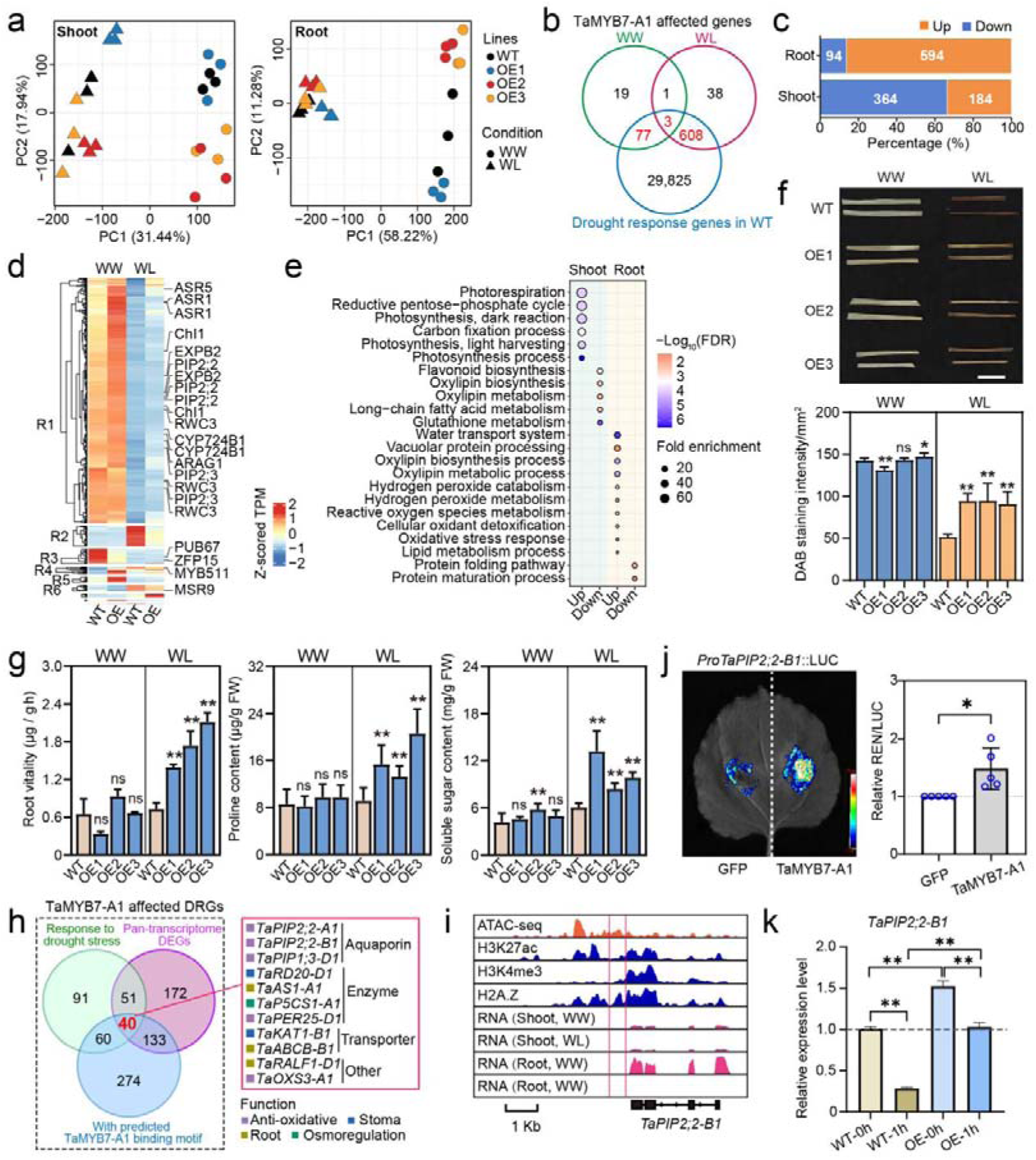
RNA-seq elucidates the molecular regulatory of TaMYB7-A1 in regulating wheat drought-resistance. **(a)** Principal component analysis of transcriptome data of shoot (left) and root (right) samples from Fielder and OE lines under different conditions. Samples of different genotype were indicated by different colors, and samples from each condition with different symbols. **(b)** Venn diagram displaying the overlap of TaMYB7-A1 affected genes under WW (green) and WL (red) conditions and drought-response genes (blue) in root. The TaMYB7-A1 affected drought response genes (DRGs) were marked in red. DEGs were identified with Log2 (FoldChange)>1 and *P*.adjust<0.05. **(c)** Proportion of drought -induced and -suppressed genes in root and shoot. **(d)** Heatmap showing the k-means clustering of TaMYB7-A1 affected DRGs in root, with reported drought-resistance genes labeled. The color intensity from blue to red represents the z-scored TPM values ranging from -2 to 2. **(e)** Enriched Gene Ontology terms for TaMYB7-A1 -activated and -repressed DRGs in shoot and root. The size of dots indicates gene ratio enriched in each term, with color representing the *P*.adjust value, which refers to the Benjamini-Hochberg procedure adjusted *p*-values of Hypergeometric test. **(f)** DAB staining of leaves in Fielder and OE lines under WW and WL conditions. Representative photographs were taken and phenotypic data investigated after air exposure for two hours (from 4 seedlings, each with three independent biological replicates; Scale bars, 1 cm). Data are means ± S.D. (*n* = 12). Student’s *t* test were used to determine the statistical significance between each OE line and Fielder. ns, no significance; *, *P* < 0.05; **, *P* < 0.01. **(g)** Comparison of the ability for osmotic regulation and resistance to oxidative damage between seedlings of Fielder and OE lines under WW and WL conditions. Data are means ± S.D. (*n* = 4 independent biological replicates). Student’s *t* test were used to determine the statistical significance between each OE line and Fielder. ns, no significance; *, *P* < 0.05; **, *P* < 0.01. **(h)** Venn diagram displaying the overlap of TaMYB7-A1 affected DRGs that with predicted TaMYB7-A1 binding motif, response to PEG-simulated drought stress, and that identified as pan-transcriptome DEGs. The 40 genes shared by all three groups were considered as potential downstream direct targets of TaMYB7-A1, and 11 genes whose homologs were known genes involved in drought-resistance or WUE were shown. **(i)** IGV showing the RNA-seq, ATAC-seq, and each histone modification across *TaPIP2;2-B1*. Red vertical lines indicate the predicted binding motif of TaMYB7-A1. **(j)** Dual-luciferase (LUC) reporter assays showing the transcriptional activation of TaMYB7-A1 on *TaPIP2;2-B1*. The relative value of LUC/REN (Renilla) was normalized with value in green fluorescent protein (GFP) set as 1. Error bars show ± S.D. of biological replicates (*n* = 5). Student’s *t*-test was used for the statistical significance. *, *P* < 0.05. **(k)** Expression levels of *TaPIP2;2-B1* in root of Fielder and OE1 under 12% PEG-simulated drought stress. RT-qPCR was used to determine their expression patterns, with *TaTublin* as the internal control. The expression levels were normalized with WT at 0 hour set to 1.0. Data are means ± S.D. of three independent biological replicates. Student’s *t* test were used to determine the statistical significance. no significant difference; **, *P* < 0.01. h, hour.

Comparing OE and WT plants, we identified equally TaMYB7-A1-affected genes under WW and WL conditions (499 *versus* 419) in shoot (**Supplementary Fig. 10a**), whereas more genes were detected under WL than WW (650 *versus* 100) in root **(Fig. 7b**). In total, 548 and 688 TaMYB7-A1-affected genes were overlapped with drought-responsive genes in shoot and root, respectively **(Fig. 7b, c, Supplementary Fig. 10a and Supplementary Table 13**). Majority of these genes were up-regulated in root, while down-regulated in shoot **(Fig. 7c and Supplementary Fig. 10c**). TaMYB7-A1-activated DRGs were enriched for GO terms related to water transport in root and photosynthesis in shoot (**Fig. 7d, e and Supplementary Fig. 10d**), aligning with the enhanced drought resistance and photosynthesis of OE lines under WL conditions (**Fig. 6**). Aquaporins (e.g., PIP2;2^74^ and RWC3^75^) and abscisic stress-ripening proteins (e.g., ASR1 and ASR5)^76^ were up-regulated in the root of OE lines, and glutamine synthetase GS2 (enhance photosynthesis under drought and salt stress)^77^ and MYB58 (produces dark green leaves)^78^ were up-regulated in shoot (**Fig. 7d, and Supplementary Fig. 10d**).

Notably, TaMYB7-A1-repressed genes in shoots were enriched for processes related to lipid peroxidation of cell membrane, whereas TaMYB7-A1-activated genes in roots were associated with oxidative stress responses (**Fig. 7d, e and Supplementary Fig. 10d**). This suggests that TaMYB7-A1 plays role in regulating antioxidant mechanisms to enhance drought tolerance in wheat. Coincidentally, key genes known for maintaining cell membrane integrity (e.g., *PIP2;2*^74^) and antioxidant defense (e.g., *ZFP182*^79^) were found (**Fig. 7d, and Supplementary Fig. 10d**). To further evaluate oxidative stress, we assessed *TaMYB7-A1* OE lines and WT plants under drought conditions. Staining of the leaf with 3,3′-diamino-bezidine (DAB) reveals reduced hydrogen peroxide (H_2_O_2_) accumulation in OE lines, indicating lower oxidative damage compared to WT plants **(Fig. 7f**). OE lines also exhibited reduced malondialdehyde (MDA) levels, a marker of lipid peroxidation, along with increased root vitality, proline, and soluble sugars contents (**Fig. 7g and Supplementary Fig. 10e**), highlighting enhanced osmotic regulation and antioxidant resilience.

To identify direct targets of TaMYB7-A1 in drought tolerance, we screened for its potential TaMYB7-A1 binding motifs in promoters of TaMYB7-A1-affected DRGS using motifs of its closet Arabidopsis orthologs (**Supplementary Fig. 10f, g**). Cross-referencing these with drought-responsive DEGs (see method for details) and pan-transcriptome DEGs, yielded 40 potential direct targets (**Fig. 7h and Supplementary Table 14**). Among these, 11 genes (27.5%) were associated with antioxidant regulation (*TaPIP2;2-A1/B1*, *TaPIP1;3-D1*, *TaOXS3-A1*, and *TaPER25-D1*)^74^, root development (*TaAS1-A1*, *TaABCB-B1*, and *TaRALF1-D1*), stomatal movement (*TaRD20-D1* and *TaKAT-B1*)^80^, and osmotic regulation (*TaP5CS-A1*)^81^ (**Fig. 7h, i and Supplementary Fig. 10h**). Luciferase (LUC) reporter assays validated that TaMYB7-A1 directly activates *TaPIP2;2-A1* and *TaPIP2;2-B1* (**Fig. 7j and Supplementary Fig. 10i**), with their expression significantly up-regulated in TaMYB7-A1-OE lines after 1-hour PEG treatment, supporting TaMYB7-A1’s role in enhancing water transport under drought (**Fig. 7k**). RT-qPCR further confirmed altered expression levels of additional target genes in OE lines, emphasising TaMYB7-A1’s broad regulatory impact (**Supplementary Fig. 10j**).

Collectively, TaMYB7-A1 enhances drought tolerance in wheat by modulating expression of genes involved in root development, osmotic regulation, stomatal movement and anti-oxidative, for instance, *TaPIP2;2*. These findings provide insights into the regulatory network underlying drought resistance and WUE in wheat.

## Discussion

With growing global populations and advancing climate change, wheat production faces mounting challenges from water scarcity, especially in arid and semi-arid regions^2,3^. Enhancing drought resistance and water-use efficiency (WUE) in wheat is essential for achieving sustainable agriculture and ensuring global food security under these climate pressures^5^. A deeper understanding of the genetic and regulatory networks governing drought resilience in wheat will empower breeding programs to develop varieties capable of withstanding future climate challenges.

### Integrated omics: advancing gene discovery and functional analysis of wheat WUE

The central dogma describes genetic information flow in biology: from DNA to RNA to protein, leading to phenotype^82^. Traditional GWAS and QTL approaches focus on individual genomic layers may over-emphasize specific loci while miss important regulatory mechanisms, gene-gene interactions, and gene-environment interactions critical for understanding polygenic and complex traits^83–85^ (**Fig. 2 and 5**).

In contrast, integrated multi-omics approaches—combining genomics, transcriptomics, proteomics, and epigenomics—provide a more comprehensive view of molecular interactions underlying complex traits^86–88^. By layering data across omics levels, we capture the dynamic, interconnected nature of biological systems and prioritize causal genes and regulatory networks associated with drought resilience (**Fig. 3 and 5**). Positioning gene expression as a key regulatory variable, this approach links genetic variations modulating expression patterns (eQTL) with downstream phenotypic effects (using WGCNA and SMR), allowing for a more precise understanding of DNA variation’s impact on observable traits under drought stress **(Fig. 5**).

Multi-omics integration also enables the identification of gene networks and stress-responsive pathways, such as those involved in drought response, osmoregulation, and phytohormone signaling in drought-tolerant wheat **(Fig. 5**). Furthermore, the discovery of pan-regulons and eQTL hotspots—clusters of loci that modulate multiple genes—adds another layer of insight **(Fig. 3**). By mapping these regulatory elements under varied conditions, we identify loci that consistently drive drought-related expression changes, crucial for breeding programs **(Fig. 3 and 4**). These insights lay the foundation for precise, base-editing crop breeding and allow for prioritization of high-potential markers in marker-assisted selection, enhancing wheat breeding for optimized WUE.

### Subgenome specific gene expression dynamics and regulatory hotspots underpin wheat drought resilience

Gene expression variation, shaped by genetic and environmental factors, drives phenotypic diversity, especially in abiotic stress responses^89,90^. Our pan-transcriptome analysis in wheat under drought stress reveals the intricate regulation of gene expression tied to drought resilience. Core genes showed high expression stability across wheat accessions, likely essential for maintaining cellular homeostasis and growth under stress, such as enzymatic activity (**Supplementary Fig. 3**). In contrast, unique genes displayed higher variability, hinting at their role in accession specific drought-induced phenotypic diversity and adaptive pathways **(Fig. 2**).

Allopolyploidy facilitates genome plasticity, allowing crops like wheat to adjust more readily to changing environmental conditions^60,89,91^. While most triads exhibited balanced expression, ensuring stable gene dosage for fundamental processes, a notable fraction showed homolog suppression or dominance **(Fig. 2**), suggesting an adaptive mechanism to fine-tune drought responses. Tissue-specific drought responses were observed: stable triads in roots were enriched for osmoregulation and oxidative stress defense, while shifts in triads in shoots were associated with photosynthesis and phytohormone pathways **(Fig. 2 and Supplementary Fig.3**). This divergence enables wheat to optimize drought resilience by tailoring responses to each tissue’s physiological needs.

Our eQTL mapping identified genomic hotspots that enhanced eGene expression levels and variability across accessions under drought conditions. These hotspots demonstrated greater detection power and phenotypic impact in GWAS, underscoring their role as regulatory elements that drive phenotypic diversity in response to stress **(Fig. 3**). Many of these hotspots aligned with known GWAS signals related to drought adaptation traits, such as pathways for “stomatal opening” and “wax biosynthesis”, regulating eGenes involved in water loss and oxidative stress response **(Fig. 3**). Notably, *cis*-eQTLs showed focused regional effects, while *trans*-eQTLs exerted broader regulatory influence, suggesting a flexible regulatory framework that supports robust drought adaptation. This combined pattern of local and distal regulation may offer the adaptability required for consistent drought resilience across varying environments.

### TaMYB7-A1: a key genetic target for enhancing drought resistance and WUE in wheat

Breeding for improved WUE in wheat has been challenging due to a limited pool of effective gene resources^92–94^. Overexpression of *TaMYB7-A1* improves drought tolerance at the seedling stage, promoting root length, volume, and surface area, enabling deeper soil moisture access and increased plant survival. At maturity, *TaMYB7-A1* overexpression positively impacts yield traits like spike number and grain size, specifically under water-limited conditions, while no compromising yield under normal conditions **(Fig. 6**).

The functional benefits of TaMYB7-A1 are likely mediated through oxidative stress reduction and osmotic regulation. By reducing hydrogen peroxide levels (as shown by DAB staining) and enhancing proline and soluble sugar levels, TaMYB7-A1 strengthens antioxidant defenses and maintains osmotic balance **(Fig. 7**). Additionally, TaMYB7-A1 lowers water loss via reduced transpiration and mitigates photosynthesis decline during drought, as reflected by higher chlorophyll content and Fv/Fm values **(Fig. 6**). Together, these mechanisms sustain photosynthesis and reduce water loss, improving WUE and biomass production per unit water.

Haplotype analysis reveals a superior variant, Hap-1, with elevated *TaMYB7-A1* expression, higher drought tolerance and WUE, making it ideal target for mark-assisted selection (MAS). The identification of *TaMYB7-A1*-associated eQTLs supports precise editing for targeted expression control, optimizing drought response while minimizing trade-offs **(Fig. 4**). These findings establish TaMYB7-A1 as a valuable foundation for advancing WUE and drought resilience in wheat under climate variability. Moreover, this study also uncovers TaMYB7-A1’s regulatory network, including regulation of *TaPIP2;2-A1* and *TaPIP2;2-B1*, providing additional resources for breeding **(Fig. 7 and Supplementary Fig. 10**).

## METHODS

### Plant materials and growth conditions

The nationwide collection of 228 wheat accessions in GWAS panel was cultivated in sealed sand plot for three weeks at Shijiazhuang, Hebei Province (37.89°N, 114.69°E) during the 2020 and 2021 (**Supplementary Table 1**). The KN9204 TILLING mutant lines were planted in the green house for drought resistance evaluation. The wheat transgenic lines were multiplied and planted in the green house under conditions of 22°C, 16h/18°C, 8h (light/dark), and 40% humidity or field of the Institute of Genetics and Developmental Biology at Changping (40.22°N, 116.23°E), Beijing. During the wheat harvest season, yield related traits were assessed, including grain length, grain width, and thousand-grain weight. The positive overexpression transgenic lines of *TaMYB7-A1* were generated and multiplied as described^73^.

Tobacco (*Nicotiana benthamiana*) was planted in a mixed soil of vermiculite, nutrient soil, and floral soil in a ratio of 1:1:1, and cultivated in greenhouse at 22L temperatures and 12h light/12h dark.

### Water stress treatment through different systems

For wheat accessions water stress experiment, three water gradients treatment was performed: 15% RSWC (well-watered, WW), 9% RSWC (moderate deficit soil water, DS1), and 6% RSWC (severe deficit soil water, DS2). Each experiment was conducted with 10 plants per pot and three replicates. Dry weight of shoot (DW.S), dry weight of root (DW.R), root surface area (RSA) and water use efficiency (WUE_p_, defined as dry biomass per unit water cost) were obtained after excluding outliers and following described as WUE-related traits.

To evaluate drought resistance at seedling stage, the method of withholding water after a period of time and then re-watering was used^37^. The germinated seeds of transgenic lines of *TaMYB7A-1* and WT (*cv.* Fielder) were transplanted into cultivation boxes (32 cm × 16 cm × 12 cm, length × width × depth) and the germinated seeds of KN9204 TILLING mutants (**Supplementary Table 12**) and WT (*cv.* KN9204) were transplanted into square boxes (8 cm × 8 cm × 8 cm, length

×width × depth), which all were filled with uniformly mixed soil (Pindstrup substrate to vermiculite in a ratio of 1:1) in the greenhouse. The soil water content was recorded by Moisture hygrometer (LY-201) every three days after water withholding, and re-watering when soil water content reduced to ∼4%. The survival rate (SR) and fresh weight (FW) were recorded 7 days later. 28 plants of each line in cultivation boxes and 36 plants of each mutant in square boxes were compared in each test.

To acquire the seedling and physiological phenotypes of transgenic lines of *TaMYB7A-1* and WT (*cv.* Fielder) under water-limited treatment, germinated seeds were transplanted into chain paper tube nursery bag (11 cm × 8 cm, diameter × depth), which were filled with uniformly mixed soil (Pindstrup substrate to vermiculite in a ratio of 1:1). When soil water content decrease to ∼20% (water-limited, WL), materials were sampled for seedling phenotype, physiological phenotype, expression analysis, RNA-seq and compared with that in water-well (WW) condition.

The WL condition also was set in field (Changping), transgenic lines of *TaMYB7A-1* and WT (*cv.* Fielder) were planted in six 1.0m-long rows under WW and WL conditions. Plants in WL condition were withheld water after flowering stage, while management of WW condition followed local cultivation practices (watering 3–4 times).

In order to further detect the expression profile of targeted genes, 1-week transgenic lines of *TaMYB7A-1* and WT seedlings were transferred to 12% PEGL6000 (m/v) stress to simulate drought treatment, samples were collected, frozen in liquid nitrogen and stored at −80°C for RT-qPCR.

### Phenotype evaluation under water-limited condition

Under WL and WW condition in green house, various seedling and physiological phenotypes of transgenic lines were examined.

After whole seedling photographed, the max length and fresh weight of shoots and roots were investigated respectively. The roots were washed cleanly and scanned by Epson Expression 13000XL, and then the root system architecture (RSA) were extract using WinRHIZO 2008 software, including total root length, total root volume and root sufferance area.

The leaf temperature of seedlings, reflecting transpiration, were photographed by InfraRed Camera R500 and analyzed using InfReCAnalyzer software. The Fv/Fm was measured by MINI-PAM-II after dark adaptation and analyzed by WinControl-3 software. The photosynthetic parameters were analyzed by LI-6400 Portable Photosynthesis system with leaf area correcting, and the WUE_i_ was conducted by instantaneous net CO_2_ absorption rate to the transpiration rate.

To evaluate reactive oxygen species (ROS) accumulation, the seedlings were stained with DAB (G4815, Solarbio) and ImageJ (v1.53k) was used to extracted gray value with the high gray value represent low ROS status.

Similarly, concerned physiological phenotypes were measured using spectrophotometry according to the instructions, including total Chl content (BC0995, Solarbio), MDA content (BC0025, Solarbio), root activity (BC5275, Solarbio), proline content (BC0295, Solarbio), and soluble sugar content (BC0035, Solarbio).

### Genotype obtaining and filtering

Genotyping of the wheat accessions was performed using Affymetrix Wheat660K SNP arrays by CapitalBio Corporation. High-quality SNP markers were filtered based on the following criteria: (1) Minor allele frequency ≥ 5%. (2) Missing genotype rate ≤ 10% across the population. (3) Heterozygosity rate ≤ 5%; (4) Uniquely mapped to the reference genome IWGSC RefSeq v1.0. A total of 323,741 SNPs (high-quality SNPs) were retained for further analysis^34^. Linkage disequilibrium (LD) decay calculation and visualization were performed using PopLDdecay (v3.42).

In addition to genotyping via Affymetrix Wheat660K SNP arrays, Whole Exome Sequencing (WES) was performed on 110 selected accessions by Tcuni (Chengdu, China). Firstly, quality control of raw reads was conducted using fastp (v0.22.0), and then BWA (v0.7.17) was used to align reads to IWGSC Refseq v1.1. Reads with ‘QUAL < 100.0, GQ < 20.0, FS > 60.0, ReadPosRankSum < -8.0, QD < 2.0, MQRankSum < -12.5, MQ < 40.0 and SOR > 3.0’ were removed. BAM files were sorted, and PCR duplicates were removed using samtools (v1.6). The GATK (v4.1.8.1) best practice workflow was used for SNP calling. The ‘HaplotypeCaller’ and ‘GenotypeGVCFs’ were used to generate and merge each sample to all sample with a raw VCF file. And bcftools (v1.9) was used to filter VCF file by ‘DP < 5 or 3 (Homozygous depth < 5, Heterozygous depth < 3), and GQ < 3’.

The final combined genotype data of 110 accessions was a combination of SNP data from the Affymetrix Wheat660K SNP arrays and WES data. This combined dataset was further filtered using bcftools (v1.9) to exclude SNPs with a missing rate > 1%, minor allele frequency < 5%, and heterozygosity rate > 50%. After filtering, a total of 697,570 high-quality SNPs were obtained and used for subsequent analysis.

### Genome-Wide Association Studies (GWAS) analysis

The high-quality SNPs of wheat accessions were used for association analysis with phenotypic data using the mixed linear model in GEMMA (v0.98.3)^95^. PCA calculated by GAPIT package (v3.4.0) was introduced as a fixed effect, the Kinship matrix was used as a random effect, and the *P*-value < 3.16E-04 was determined to be the suggestive threshold for declaring significant association. Manhattan plots and quantile-quantile plots were generated using CMplot package (v4.5.1), and LD plots were calculated and displayed using LDBlockShow (v1.40). The phenotypic variation explained was calculated based on the effect size, standard error, and minor allele frequency according to the output parameters.

### RNA extraction, sequencing and population-transcriptome analysis

Total RNA (110 selected wheat accessions, three water gradients stress, two tissues) was extracted using the Quick RNA Isolation Kit (2407#22D, HUAYUEYANG BIOTECHNOLOGY CO., LTD) according to the manufacturer’s instructions. Next, the 3’ RNA-seq libraries were constructed as described with slight modification^45^. In order, fragmenting RNA (∼2 μg), synthesizing the first-strand cDNA and the double-stranded DNA, performing end repair, dA-tailing and adapter ligation, finally PCR amplification. The libraries were sequenced on NovaSeq platform (Illumina) for 150 bp paired-end reads. Raw data were filtered to remove adapters and low-quality bases using fastp (v0.22.0) and clean data were mapped to the IWGSC Refseq v1.0 reference genome using hisat2 (v2.2.1), resulting 638 high-quality RNA-seq samples for subsequent analysis (**Supplementary Table 3**). Gene count matrixs were generated by featureCounts (v2.0.3), and the expression level [Counts Per Million (CPM)] was calculated for each gene by edgeR package (v3.42.4) in R (v4.2.2).

According to data distribution characteristics, the genes with decile expression levels more than 1 (CPM > 1) among the population were considered as expressed genes, and also were transformed to a normal distribution with the R function ‘qqnorm’ (v3.6.1) for following eQTL identification. Differentially expressed genes (DEG) were identified using the edgeR package (v3.42.4) with the criteria (|log_2_FC| ≥ 1, *P*-value ≤ 0.05), and GFOLD (v1.1.4) (|log_2_FC| ≥ 1) with occurred at least 15% of accessions under any kind of water stress condition, respectively^49^. K-means cluster analysis was performed in R (v4.2.2), and gene expression heatmap was displayed using ComplexHeatmap package (v2.14.0). The definition and visualization of homoeolog expression bias were performed as previous described by rdist (v0.0.5) and ggtern (v3.5.0), respectively^60^.

### Expression quantitative trait locus (eQTL) and hotspot analysis

The mixed linear model implemented in MatrixEQTL package (v2.3)^96^ and GEMMA (v0.98.3) which considering the population structure, genetic relatedness and the estimated confounding factors were simultaneously used for association analysis between the combined genotype and the normalized expression profile of expressed gene.

The significantly associated SNPs (*P*-value < 4.11E-07, determined by 0.05/effective SNP number, which calculated using the Genetic Type I error calculator software) identified by MatrixEQTL and GEMMA were intersected^97^. A revised two-step method was employed to address associations involving multiple SNPs for a single gene expression and to define eQTL region: (1) all of the associated SNPs were grouped into one cluster if the distance between two consecutive SNPs was less than 10 kb, and the clusters with at least three significantly associated SNPs were considered as candidate eQTLs represented by their lead SNP which had the most significant *P*-value. (2) Candidate eQTLs in linkage disequilibrium (*r²* > 0.1) with other more significant candidate eQTLs for the same expression trait were identified as LD-driven associations and subsequently removed. According to the LD decay distance of combined genotype, the cutoff of distance between lead SNP and associated gene (named eGene) was set 2.4 Mb, identifying *cis*-eQTLs and *trans*-eQTLs.

eQTL hotspot was considered a genomic region which regulate the expression of a certain number of genes, and the gene number threshold was calculated and confirmed by 1,000 random permutations (*P*-value < 0.01). Circos plot was displayed by TBtools (v1.6).

### Weighted Correlation Network Analysis (WGCNA)

The Weighted Correlation Network Analysis (WGCNA), based on the concept of scale-free networks, was applied to identify gene modules which had high correlation in gene expression among population accessions^98^. The steps were followed below with WGCNA package (v1.72.5) in R: Removing outlier samples and genes, calculating similarity matrix, determining soft threshold (SFT, β power), converting similarity matrix to adjacency matrix, converting adjacency matrix to topological overlap matrix (TOM), constructing modules by one-step method (minModuleSize = 50, mergeCutHeight = 0.25), analyzing correlation between modules and phenotype traits, excavating interested modules and genes, and visualizing co-expression network by Cytoscape (v3.7.2).

### Summary data-based Mendelian randomization (SMR) analysis

After prepared data formatting, summary data-based Mendelian randomization analysis (SMR, v1.3.1)^99^ was applied to evaluate the association between gene expression and trait variation under different environment. The summary-level statistic was analyzed using the 10,000 Kb and 5,000 Kb windows for *cis*-eQTLs and *trans*-eQTLs, respectively. Heterogeneity in dependent instruments outlier test was applied to separate pleiotropy which association due to linkage. SMR associations were declared significant if *P*-value < 0.01 and the HEIDI-test *P*-value > 0.05^62,100^.

### Construction of co-localization network and transcriptional regulation network

The GWAS bin was defined as the candidate interval which merged and extended 3 Mb (according to LD decay distance) by a single SNP passed the threshold in GWAS analysis. For the significant causal genes in SMR analysis, the SNPs controlling gene expression were identified by combining with eQTL analysis, and the SNPs were required to be located in significantly associated GWAS bin. Each trait was analyzed independently and finally a physical location-based colocation network was built.

For candidate genes locked by co-localization and DEG of population-transcriptome, potential transcriptional regulatory networks was constructed based on the correlation of gene expression levels (*P*-value < 0.05) and the motif binding target genes by TF (*P*-value < 1E-04 in FIMO scanning) in different tissues. It is stated that the matched motif used was in the PlantTFDB database (https://planttfdb.gao-lab.org/).

### Expression analysis

Total RNA were extracted using Quick RNA Isolation Kit (2407#22D, HUAYUEYANG BIOTECHNOLOGY CO., LTD). First-strand complementary DNA (cDNA) was synthesized from 2 μg of DNase I-treated total RNA using the FastKing RT kit (KR116, TIANGEN BIOTECH CO., LTD). RT-qPCR was performed using the ChamQ Universal SYBR qPCR Master Mix (Q711-03, Vazyme) by QuantStudio^TM^ 5 (Applied Biosystems, Thermo Fisher Scientific). The relative expression of interested genes was normalized to the Tubulin gene (*TaTubulin*, *TraesCS1D02G353100*) via the 2^−ΔΔCT^ method, and RT-qPCR primers used were detailed in **Supplementary Table 15**.

### RNA-seq of transgenic lines of *TaMYB7-A1* and WT

After adapters removing, low reads filtering, reference genome mapping (IWGSC Refseq v1.1) and counts calculating, transcripts per million (TPM) was normalized. The Principal Component Analysis (PCA) and Heatmap of cluster were analyzed and visualized by FactoMineR (v2.11) and ComplexHeatmap package (2.14.0) respectively. DEGs were identified using DESeq2 (v1.34.0) with the threshold of “Log_2_(FoldChange) > 1 and *P*.adjust < 0.05”.

The evolutionary tree was constructed based on sequence similarity in MEGA 7 to select the matched motif to scan latent downstream gene targeted by TaMYB7-A1, and FIMO (v 5.5.5) was used to scan motif in the 2 Kb promoter sequences of TaMYB7-A1 regulated genes (with default parameters *P*-value = 1E-04). On this basis, downstream candidate genes were also considered whether DEG of population-transcriptome and whether response to drought (available at SRP045409 and SRP068165) to further narrow the target^101^.

### Luciferase reporter assay

The promoter of *TaPIP2;2-A1* and *TaPIP2;2-B1* (∼2 Kb) were amplified from Chinese Spring (CS), and fused into CP461-LUC vector to construct the *ProTaPIP2;2-A1::LUC* and *ProTaPIP2;2-B1::LUC* reporter vectors. The coding sequences (CDS) of *TaMYB7-A1* from CS was cloned into the pSUPER-GFP vector (PRI101) as effectors. These effector and reporter plasmids were transformed into *Agrobacterium* GV3101 and injected into *Nicotiana benthamiana* leaves (at 6-8 leaf stage) in different combinations. After cultivated in greenhouse at 22°C temperatures and 12h light/12h dark for 2-3 days, the Firefly luciferase (LUC) and Renilla luciferase (REN) activities were measured using Dual-Glo^®^ Luciferase Assay System (E2940, Promega Biotech Co., Ltd). Fluorescence signals were quantified using the dualLLUC assay reagent (MOLECULAR DEVICES, SpectraMax iD3), and relative LUC activity was determined by calculating the ratio of LUC to REN. Primers for vector construction were listed in **Supplementary Table 15**.

### Statistics and data visualization

Unless specified, R (https://cran.r-project.org/v4.2.2) and GraphPad Prism (v8.0) were used to compute statistics and generate plots. Pearson correlation analysis was used for the correlation between two groups of data (**Fig. 1c, 1f, 1j, 2i, 4a, 4j, 5g, and Supplementary Fig. 1a, 3f, 6b, 6e, 9**), and corresponding *P*-values were calculated by the Pearson correlation coefficient analysis (**Fig. 1f, 1j, 2i, 4a, 4j and Supplementary Fig. 1a, 3f, 6b, 6e**). The Fisher’s exact test were used for enrichment analysis (**Fig. 2h, 2m, 3i, 4b, 7e and Supplementary Fig. 3c, 3e, 3h, 6f, 10b**). For comparisons between two groups of data conforming to a normal distribution, Student’s two tailed *t*-test was used (**Fig. 1a, 5i, 6c-h, 6j, 6k, 7f, 7g, 7j, 7k and Supplementary Fig. 8a-c, 10e, 10i, 10j**). For two-group comparison of data that did not conforming to a normal distribution, the Wilcoxon rank-sum test was used **(Fig. 1h, 1i, 2b, 2c, 2e, 2f, 3e, 3j, 3k, 4d, 4i, 5d and Supplementary Fig. 3b, 5h, 5i, 6h)**. Tukey’s test was used for multiple comparisons **(Fig. 1e, 1j, 6b, and Supplementary Fig. 1c, 2g)**.

## Data and code availability

The raw sequence data in this study was deposited in the Genome Sequence Archive database in the National Genomics Data Center (https://bigd.big.ac.cn) under accession number PRJCA032349. Source code for analysis is available at https://github.com/yx-zhou/wheat-WUE.

## Author contributions

J.X. and D.-Z. W. designed and supervised the research, Y.-X. Z., P.Z. and X.-M. L. performed the omics data analysis, Y.C. performed the RNA-seq analysis of *TaMYB7-A1* overexpression lines. Y.-Z. Q., B.-D. D., D.-Z.W., and H.W. did the evaluation of wheat accessions WUE-related traits. Y.-X. Z. did most of the experiments, H.W. generated the overexpression lines of *TaMYB7-A1*, evaluated the yield related trait in field, performed luciferase reporter assay. Y.-M.Y. helped on the 3’ RNA-seq library construction. Y.-X. Z. and D.-Z. W. prepared all the figures. D.-Z. W., J. X. and Y.-X. Z. wrote the manuscript. X.-L. L., S.-B. X. and B.-D.D. provided comments on the manuscript. All authors discussed the results and commented on the manuscript.

## Acknowledgements

We thank Prof. Zhaorong Hu from China Agricultural University for the advice and help in drought resistance evaluation of transgenic materials.

## SUPPORTING INFORMATION

**Supplemental Table 1. The wheat accessions in GWAS panel and population-transcriptome sequencing**

**Supplemental Table 2. Identified QTLs for WUE-related traits in the GWAS**

**Supplemental Table 3. Summary of reads mapping of population-transcriptome sequencing**

**Supplemental Table 4. Identified differentially expressed genes in the population-transcriptome**

**Supplemental Table 5. The enriched Gene Ontology terms for population-transcriptome gene clusters**

**Supplemental Table 6. The state transition of triads across water conditions**

**Supplemental Table 7. Identification and classification of eQTLs in each environment**

**Supplemental Table 8. Genes in core modules of WGCNA**

**Supplemental Table 9. The enriched transcription factors family in the promoter of genes in red and darkRed modules**

**Supplemental Table 10. Genes significantly associated with traits in SMR analysis**

**Supplemental Table 11. The co-localization of combing GWAS, eQTL and SMR**

**Supplemental Table 12. The KN9204 TILLING mutant lines used for candidate genes**

**Supplemental Table 13. The TaMYB7-A1 affected drought response genes identified in OE lines**

**Supplemental Table 14. The identification of potential direct targets for TaMYB7-A1**

**Supplemental Table 15. Primers used in this study**

## FIGURES LEGENDS

**Supplementary Fig. 1.**
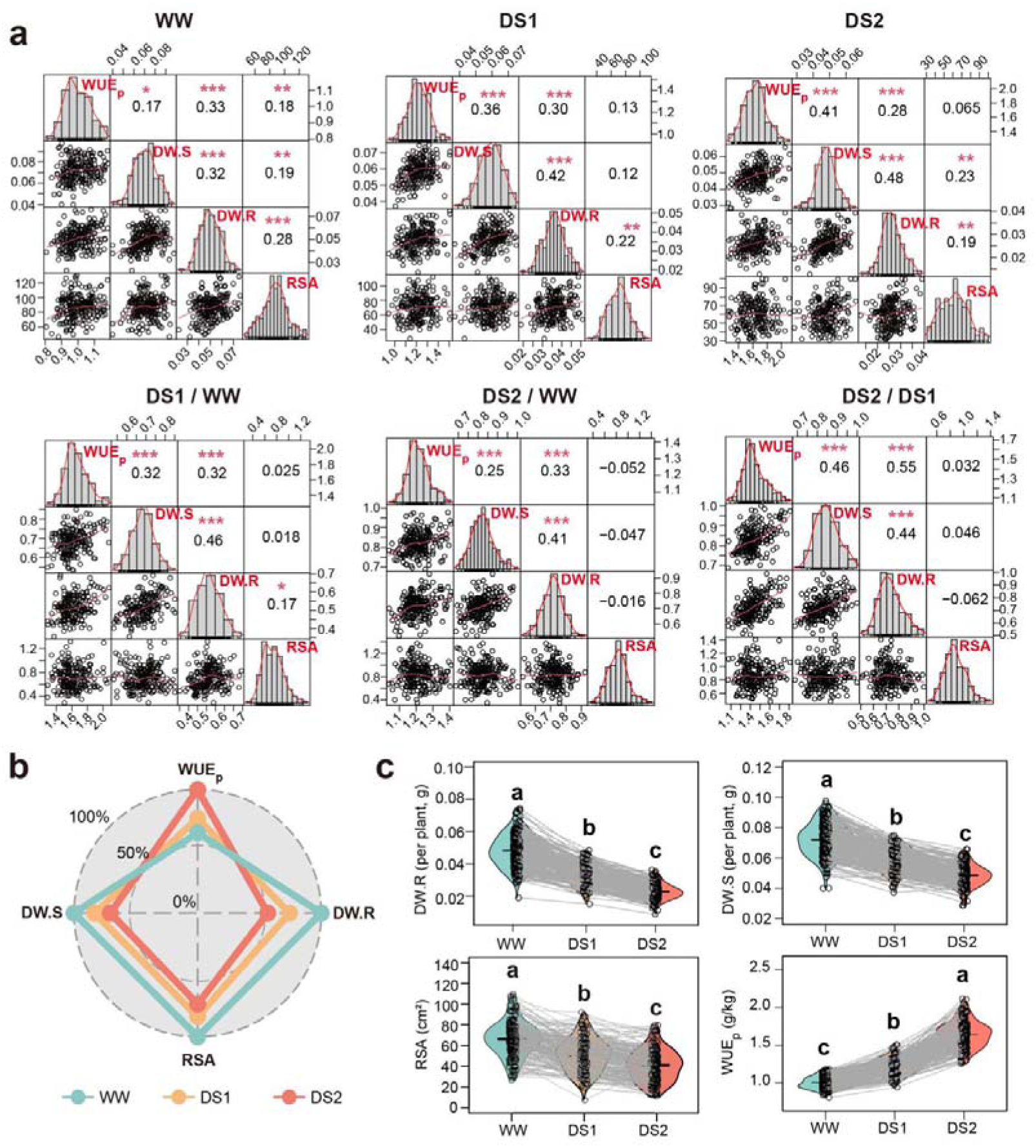
Trait performance and correlation analysis across different water regimes. **(a)** Distribution and pairwise correlation of DW.R, DW.S, WUE_p_ and RSA among environments (WW, DS1, DS2, DS1/WW, DS2/WW and DS2/DS1). The frequency distribution of each trait was shown in the histogram at the diagonal cells. The X-Y scatter plot showing the correlation between traits at the lower-triangle panel, while the corresponding Pearson’s coefficients between each trait were showed at upper-triangle panel. *, *P* ≤ 0.05; **, *P* ≤ 0.01; ***, *P* < 0.001. **(b)** Data for each trait are presented relative to the maximum value across all three conditions, with the population average shown. **(c)** Differences in drought vulnerability among wheat varieties. Each accession line is plotted as a point, and that of different conditions linked with a grey line. Tukey’s HSD multiple comparison test was used to compare the trait values among each conditions, with different letters indicate a statistically significant difference at *P* < 0.05.

**Supplementary Fig. 2.**
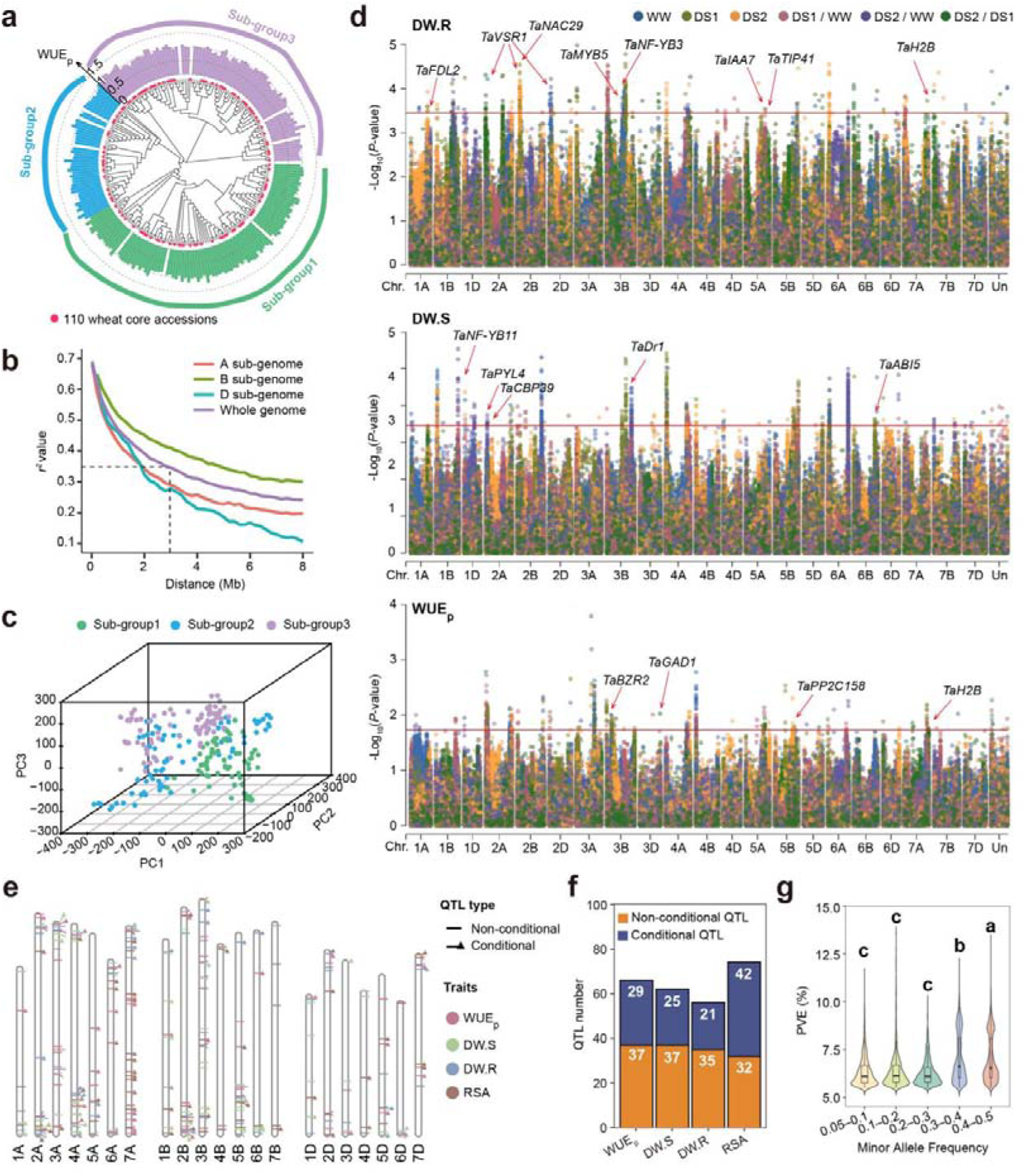
Population structure and GWAS analysis. **(a)** Phylogenetic trees were constructed using the Neighbor-Joining method, dividing the population into three sub-groups. WUE_p_ of each accession under WW conditions is plotted in a histogram surrounding the phylogenetic trees, with accessions from different sub-groups shown in different colors. The 110 accessions forming the core set are marked with red circles. Tree scale = 0.1. **(b)** Decay of linkage disequilibrium (LD) in the whole genome and three sub-genomes of the 230 wheat accessions. The physical distance when LDs of the whole genomes decayed to half of the initial value was estimated as 3 Mb. **(c)** Principal component analysis (PCA) showing the population structure of the 228 wheat accessions used in GWAS panel. The accessions in different subgroups were shown in different color dots. **(d)** Manhattan plots illustrating the Llog_10_(*P*) values for DW.R, DW.S and WUE_p_ across multiple environments. Horizontal lines indicate the genome-wide significant threshold of Llog_10_(*P*) = 3.50. Dots of different colors represent different environments. Wheat orthologs of reported drought-resistance genes located in the linkage disequilibrium (LD) region were labeled. **(e)** The chromosomal locations of conditional and non-conditional QTLs for DW.R, DW.S, WUE_p_, and RSA are indicated by different symbols. QTLs for each trait are represented in distinct colors, with lines marking the positions where the corresponding QTLs are located. **(f)** The number of conditional and non-conditional QTLs for DW.R, DW.S, WUE_p_, and RSA. **(g)** Comparison of the marker *R^2^* values in GWAS for SNPs with different minor allele frequency. SNPs passed the significance threshold of Llog_10_(*P*) = 3.50 were retained, and data from different traits and conditions were polled together. Tukey’s HSD multiple comparison test was used to compare the □log_10_(*P*) value between each group, with different letters indicate a statistically significant difference at *P* < 0.05.

**Supplementary Fig. 3.**
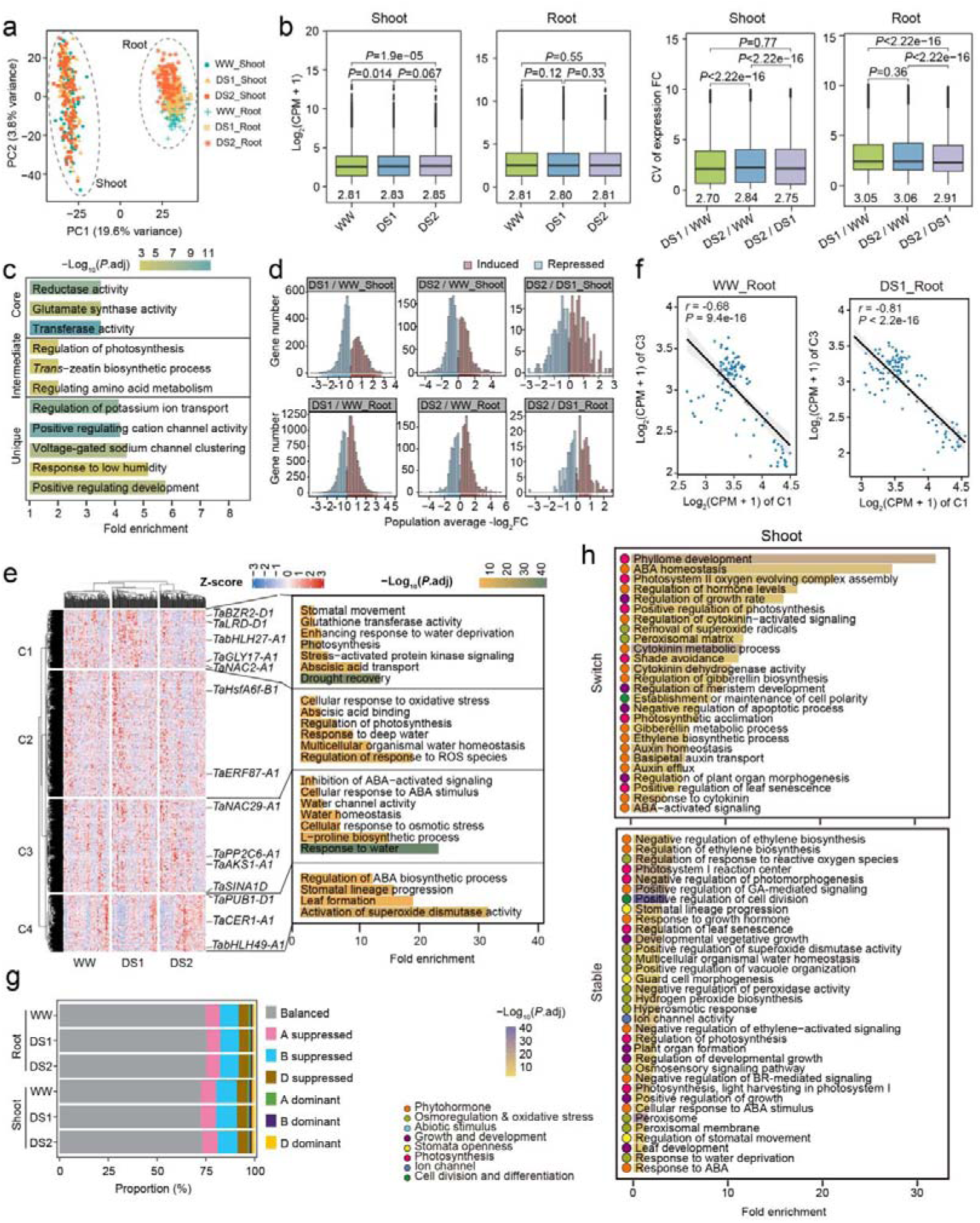
The subgenome expression bias of wheat encountering water scarcity. **(a)** Principal component analysis (PCA) of population-level transcriptome data of root and shoot samples under different conditions. Dashed ovals separate the root samples and the shoot samples. **(b)** Comparison of the expression level (left), and CV for expression foldchange (right) among expressed genes in shoot and root under different conditions. In the box plot, borders represent the first and third quartiles, center line denotes median, and whiskers extend to 1.5 times the interquartile range beyond the quartiles. Wilcoxon rank sum test was used to determine the statistical significance between two groups. **(c)** Enriched Gene Ontology terms for core, unique, and intermediate genes in root. The length of shade indicates gene ratio enriched in each term, with color representing the -log_10_(*P*.adjust). Fisher’s exact test were used for enrichment analysis. Circles in the left indicate the functional group of GO terms. **(d)** Distribution of DEGs identified in each comparable group across different population average -log_2_FC values. For each gene, the -log_2_FC values in all accessions were averaged. The induced- and repressed-genes in each comparable group were marked in brown and blue, respectively. **(e)** Heatmap showing the k-means clustering of DEGs in shoot, with reported drought-resistance genes labeled, and the top enriched Gene ontology terms shown on the right. Fisher’s exact test were used for enrichment analysis. **(f)** The scatterplots showing the correlation of population-level average gene expression between C1 and C3 in root under WW and DS1 conditions. Each dot represents an accession. The black line represents the regression trend calculated by the general linear model, and the 95% confidence intervals were indicated with gray shade. The *r* value and *P* values were determined by Pearson correlation coefficient analysis. **(g)** Proportion of triads in each category of homoeolog expression bias across the tissue and water conditions. All the triads with at least one homoeolog with CPM > 0.5 were used. **(h)** Sankey diagram showing the state switching of triads for expressed genes under different water conditions. All the triads with at least one homoeolog with CPM > 0.5 were used. **(i)** Enriched Gene Ontology terms for switch (upper panel) and stable (lower panel) triads in shoot. The length of shade indicates gene ratio enriched in each term, with color representing the - log_10_(*P*.adjust). Fisher’s exact test were used for enrichment analysis.

**Supplementary Fig. 4.**
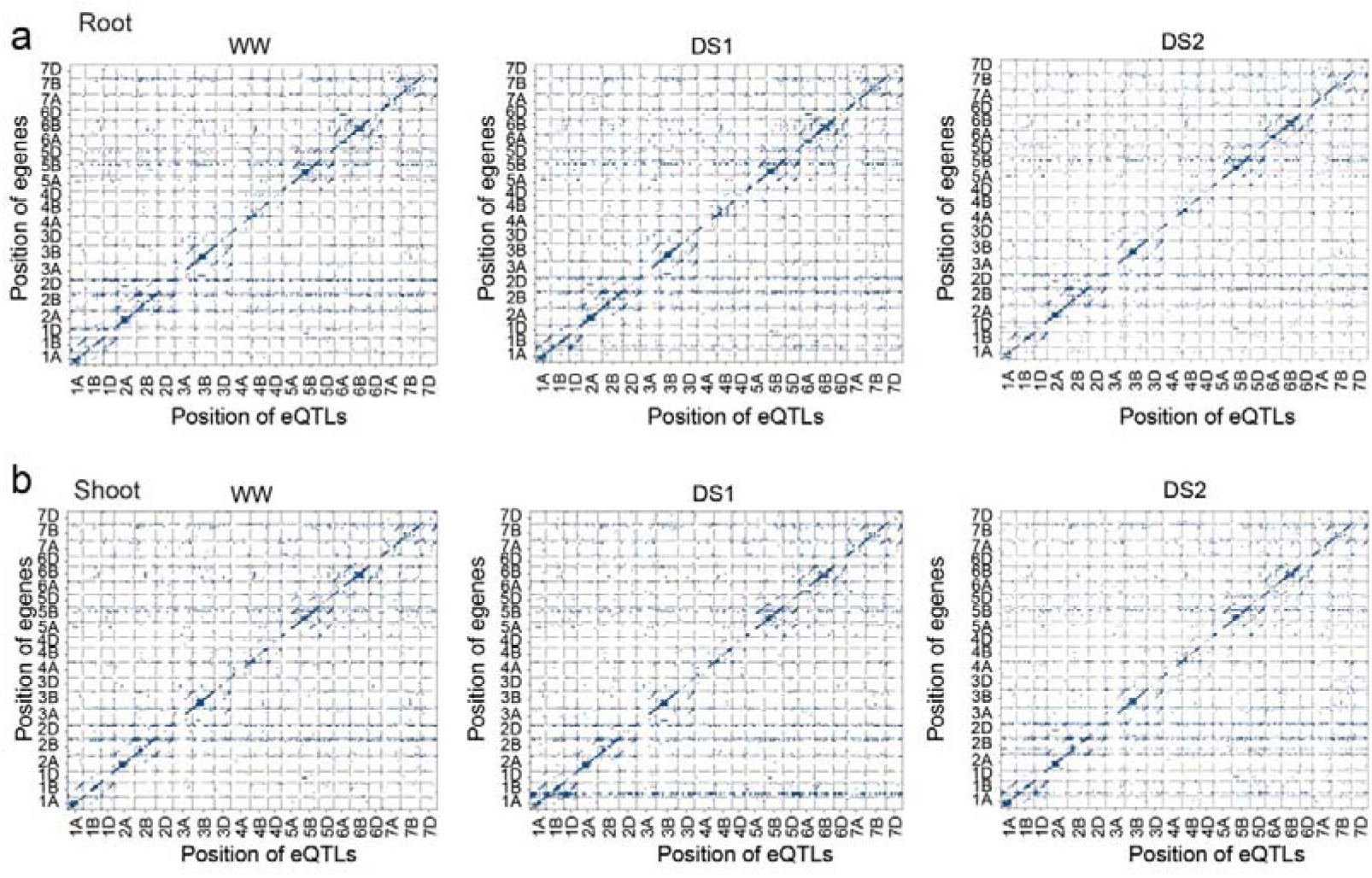
Location of eQTL relative to positions of their regulatory eGenes on 21 chromosomes in root (a) and shoot (b) X-axis, SNP position on each chromosome; Y--axis, eGene position on each chromosome.

**Supplementary Fig. 5.**
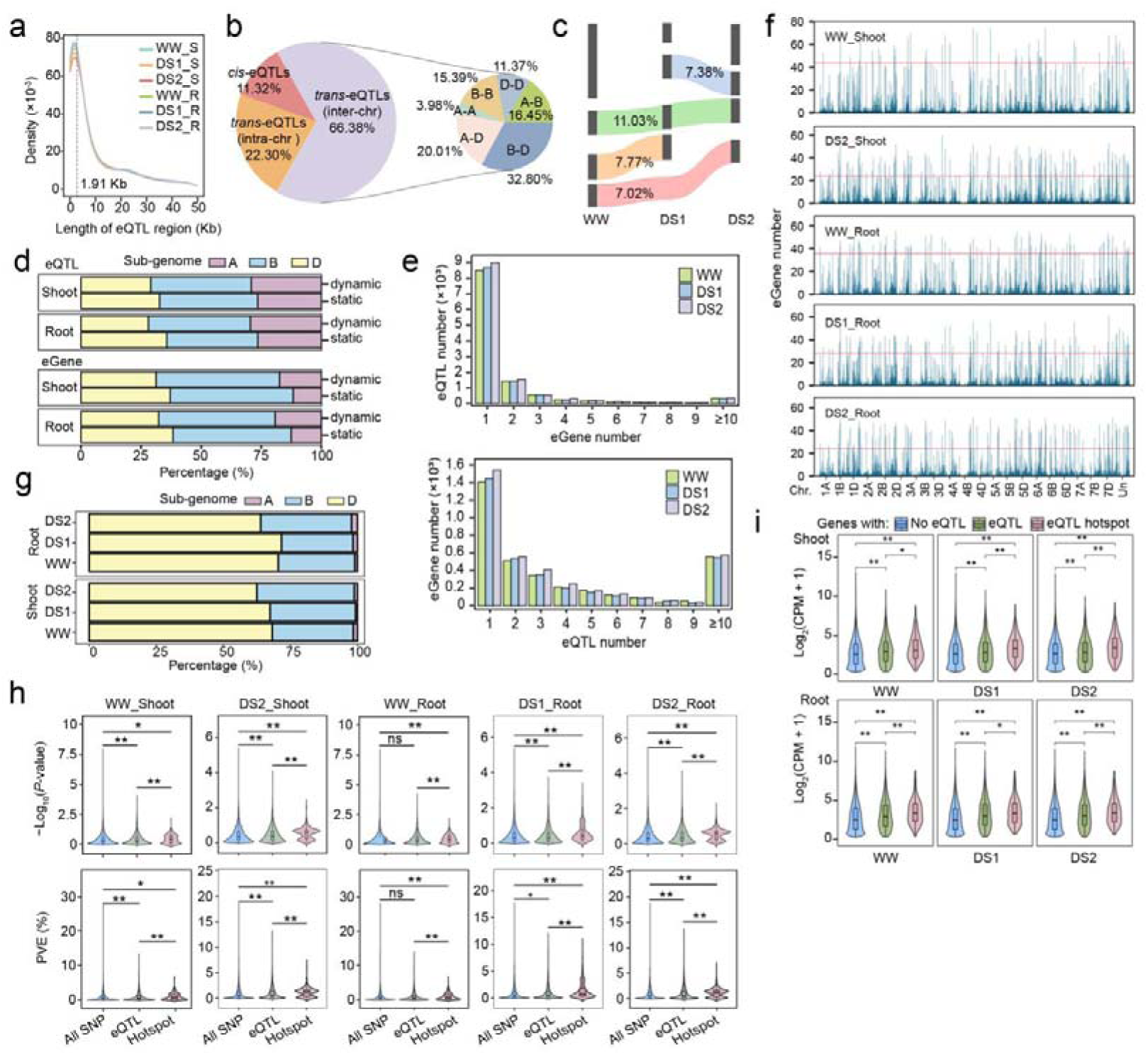
Expression diversity shapes the phenotypic resilience of wheat to drought stress. **(a)** Density distribution of eQTL region length. The dashed vertical line indicates the average eGene region length by pooling eQTL regions of each tissue and water condition. **(b)** Proportion distribution of *cis*- and *trans*- eQTLs and the subgenomic interactions involved in *trans*-eQTL regulation in shoot tissue. The inter-eQTLs indicates that eQTL and eGenes located in the same chromosome, and intra-eQTLs means that the eQTL and its eGenes are on different chromosomes. **(c)** Sankey diagram showing the sharing and specificity of eQTLs in shoot among water conditions. The numbers indicated proportion of eQTLs of each group relative to the total eQTL number. **(d)** eQTLs and eGenes distribution of each category on three subgenomes. The eQTL regulating more than one genes and the eGene regulated by more than one eQTL was counted only once **(e)** Distribution of eQTLs with each number of target genes (upper panel), and eGenes with each number of eQTLs (lower panel) in shoot across different water conditions. **(f)** Number of eGenes regulated by distal eQTLs identified in different conditions. The horizontal dashed lines indicate the threshold of eGene numbers regulated by eQTL hotspot regions. **(g)** Distribution of hotspot eQTLs regulating genes on three subgenomes across the tissue and water conditions. The eGene regulated by more than one eQTL was counted only once. **(h)** Comparison of -log_10_(*P*-value) and phenotypic variation of SNPs explained in the GWAS of WUE_p_ among all GWAS SNPs, non-hotspot eQTLs and hotspot eQTLs. The non-hotspot eQTLs and hotspot eQTLs in each condition were used. Wilcoxon rank sum test was used to determine significant differences. ns, no significance; *, *P* < 0.05; **, *P* < 0.01. **(i)** Expression levels of genes with no eQTL, non-hotspot eQTLs and hotspot eQTLs. Wilcoxon rank sum test was used to determine significant differences. *, *P* < 0.05; **, *P* < 0.01.

**Supplementary Fig. 6.**
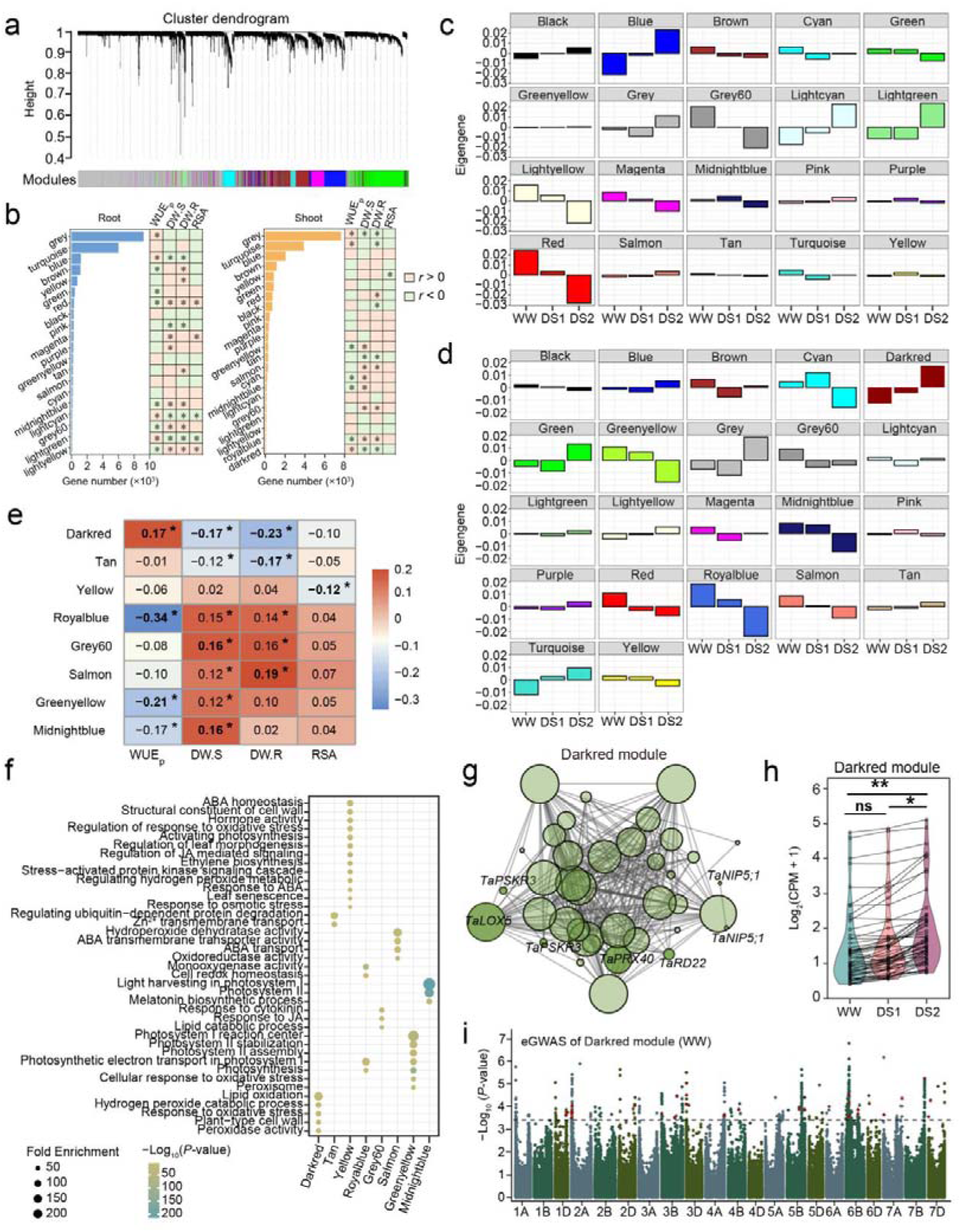
Dissecting co-expression modules underlying phenotypic diversity for drought-resistance. **(a)** Hierarchical cluster dendrogram showing co-expression modules in shoot. The co-expressed modules were identified by weighted gene co-expression network analysis and each module was indicated with a branch, and each gene was indicated with a leaf with ordinate showing the clustering distance of each gene. **(b)** Heatmaps showing the gene number of co-expression modules and the correlation between trait and the modules in root (left) and shoot (right). The modules whose expression positively correlated (*r* > 0) and negatively correlated (*r* < 0) with each trait were marked in orange and green, respectively. The modules significantly correlated with each trait (*P* < 0.05) were indicated by asterisks. The *r* value and *P* values were determined by Pearson correlation coefficient analysis. (**c, d**) Bar graph showing the expression of the eigengenes for each module in root (c) and shoot (d) tissues under different water conditions. The expression of the eigengenes in each module were normalized by setting average eigengene expression across all samples (all accession from three water conditions) as zero. Higher or lower expression of the eigengene under water conditiona were indicated in the vertical axis direction in each bar graph. **(e)** Heatmap showing top three correlated modules of each trait in shoot. Different colors showing the Pearson correlation coefficient, with values displayed in the square and asterisks indicated the significant correlation with *P* < 0.05. **(f)** Enriched Gene Ontology (GO) terms for top correlated modules in (e). The size of dots indicates gene ratio enriched in each term, with color representing the *P*.adjust value. Fisher’s exact test were used for enrichment analysis. **(g)** Co-expression gene network in the Darkred module in shoot. The size of the cycles indicates number of linked genes in the module. **(h)** Comparison of the expression level of Darkred module in shoot among three water conditions. Each gene in the module is plotted as a dot, and their expression levels under different water conditions are linked with a gray line. Wilcoxon rank sum test was used to determine significant differences. ns, no significance; *, *P* < 0.05; **, *P* < 0.01. **(i)** Manhattan plots illustrating the eGWAS □log_10_(*P*-value) values using the expression level of the Darkred module as a trait. Horizontal dashed line indicates the genome-wide significant threshold of □log_10_(*P*-value) = 3.50. The SNPs identified as the hotspot eQTL in root were marked in red.

**Supplementary Fig. 7.**
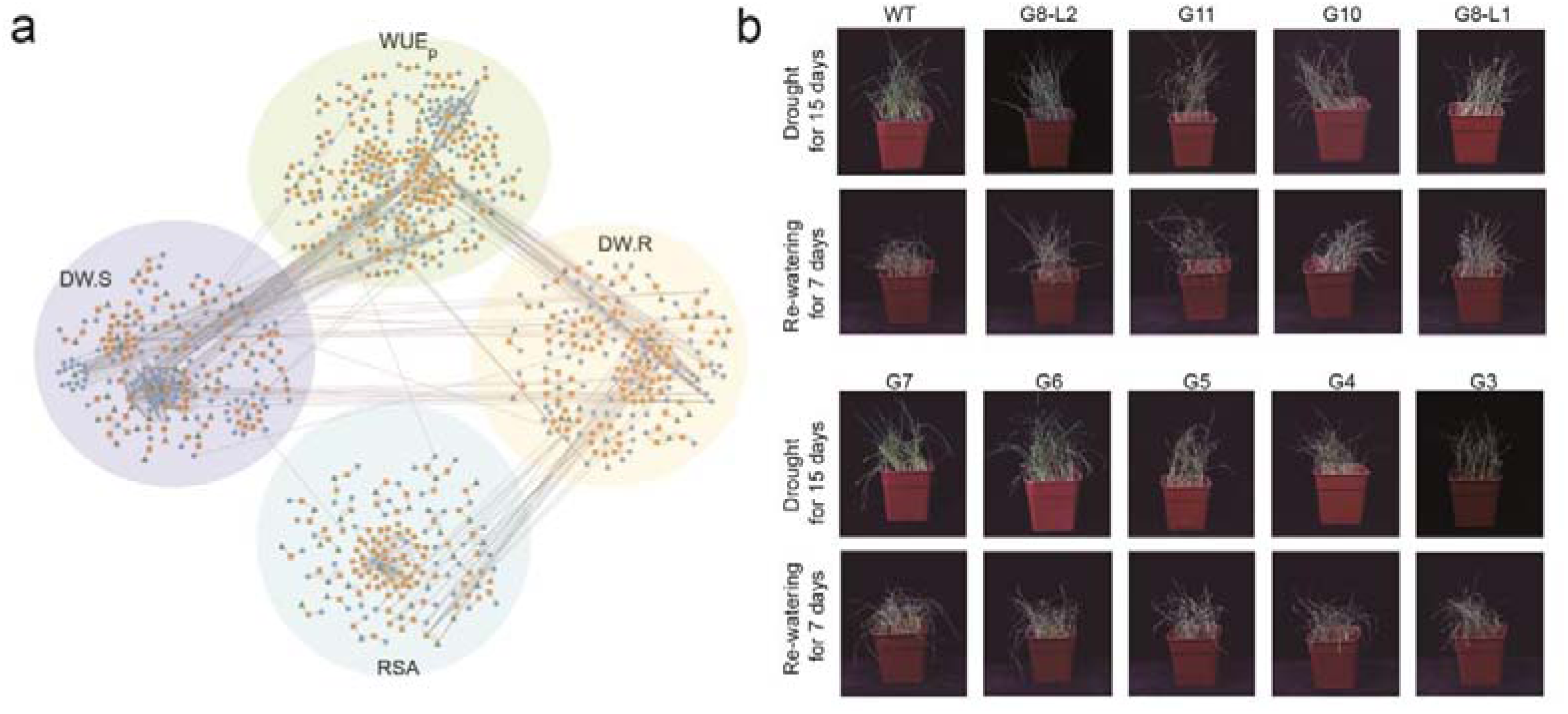
Assessment of genetic effects for candidate genes. **(a)** The genetic network of WUE_p_, DW.R, DW.S, and RSA constructed by integrating GWAS signal, eQTL and SMR genes. The GWAS signal, eQTL and SMR genes were indicated with different symbols, and the line between SMR gene and eQTL indicates a regulatory relationship, and lines between GWAS signals and SMR genes or eQTLs indicate co-localization of LD regions. The genetic network of WUE_p_, DW.R, DW.S, and RSA were marked in shade with different colors, and lines between genetic networks of different traits indicate co-localization GWAS signal with SMR genes or eQTLs in other traits. **(b)** Representative photographs showing the phenotype of wild-type KN9204 (WT) and several loss-of-function mutant lines under drought stress. Photos (each biological replicate with 36 seedlings) were taken after water-holding for 15-days (upper panel) and then water recovery for 7-days (lower panel).

**Supplementary Fig. 8.**
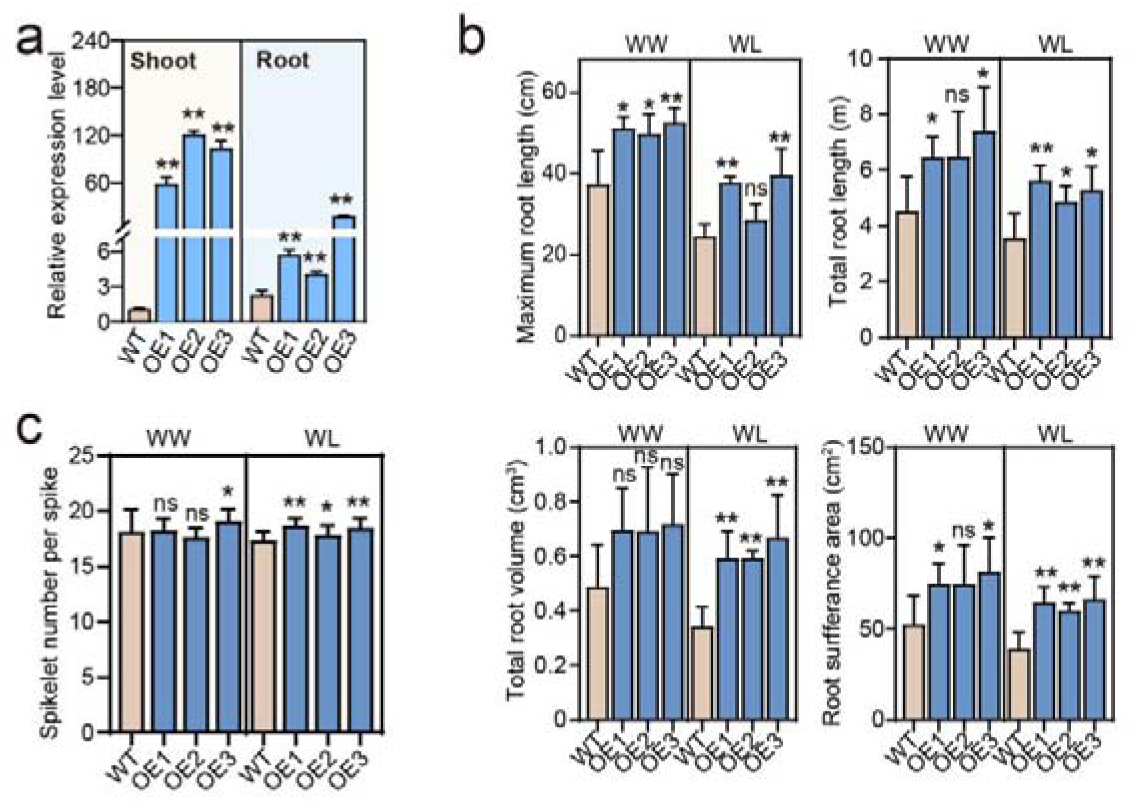
TaMYB7-A1 enhances osmotic regulation and resistance to oxidative damage under drought conditions. **(a)** RT-qPCR verifying the overexpression of *TaMYB7-A1* in shoot and root of OE lines. *TaTublin* was used as the internal control. Data are means ± S.D. of three independent biological replicates. Student’s *t* test were used to determine the statistical significance. **, *P* < 0.01. **(b)** Comparison of the root morphology between seedlings of Fielder and OE lines under WW and WL conditions. Data are means ± S.D of five independent biological replicates.. Student’s *t* test were used to determine the statistical significance between each OE line and Fielder. ns, no significance; *, *P* < 0.05; **, *P* < 0.01. **(c)** Comparison of spikelet number per spike between Fielder and OE lines under WW and WL conditions in field. Data are means ± S.D. (*n* = 40 independent biological replicates). Student’s *t* test were used to determine the statistical significance between each OE line and Fielder. ns, no significance; *, *P* < 0.05; **, *P* < 0.01.

**Supplementary Fig. 9.**
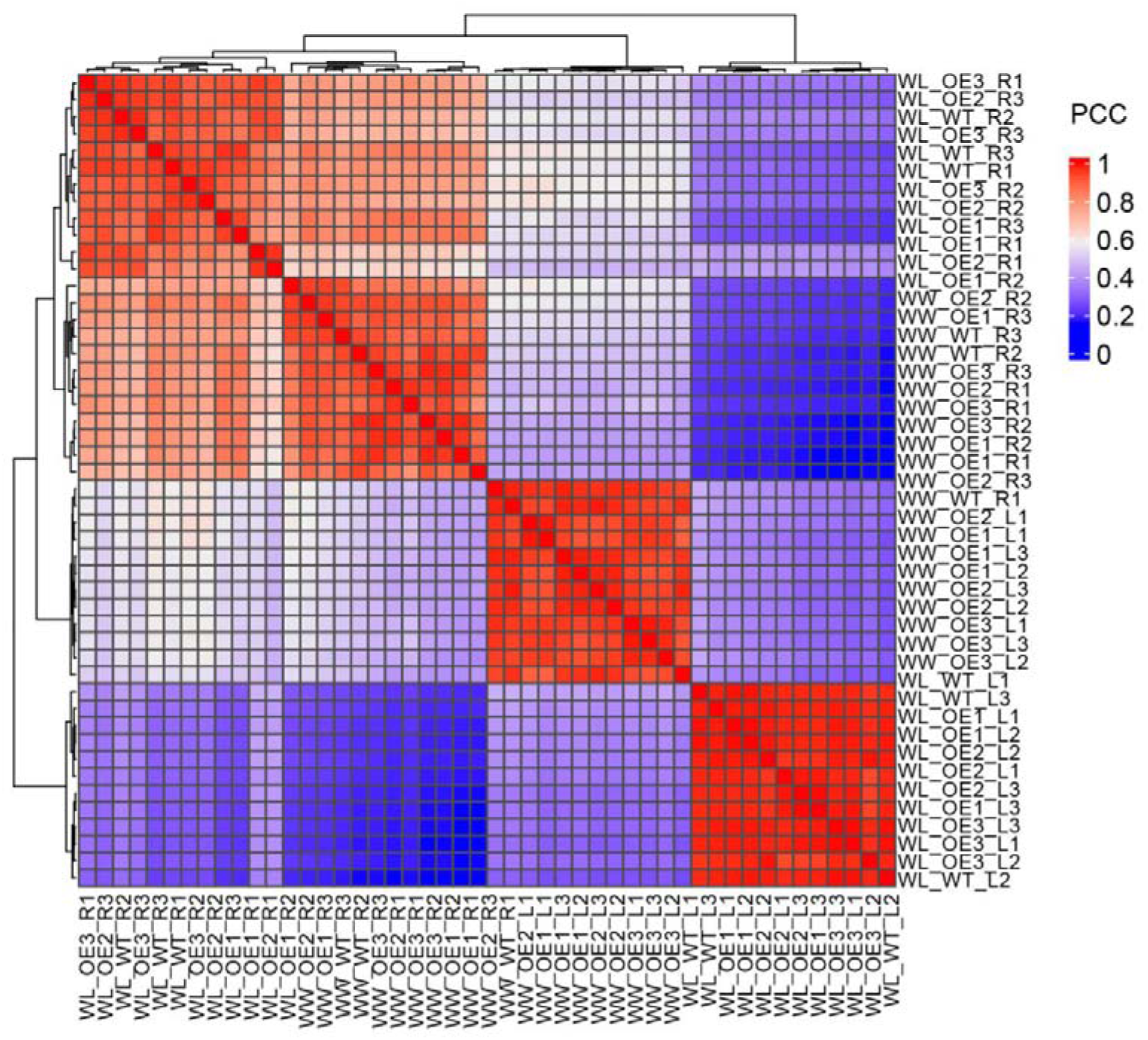
Pair-wise correlation map among RNA-seq samples from Fielder and OE lines under WW and WL conditions. The sample were naming in the format of “Condition_Genotype_Tissue & technical replicates”. L, Shoot; R, Root. The color of squares indicate the Pearson correlation coefficient (PCC) scores with color intensity from blue to red represents PCC values ranging from 0 to 1.

**Supplementary Fig. 10.**
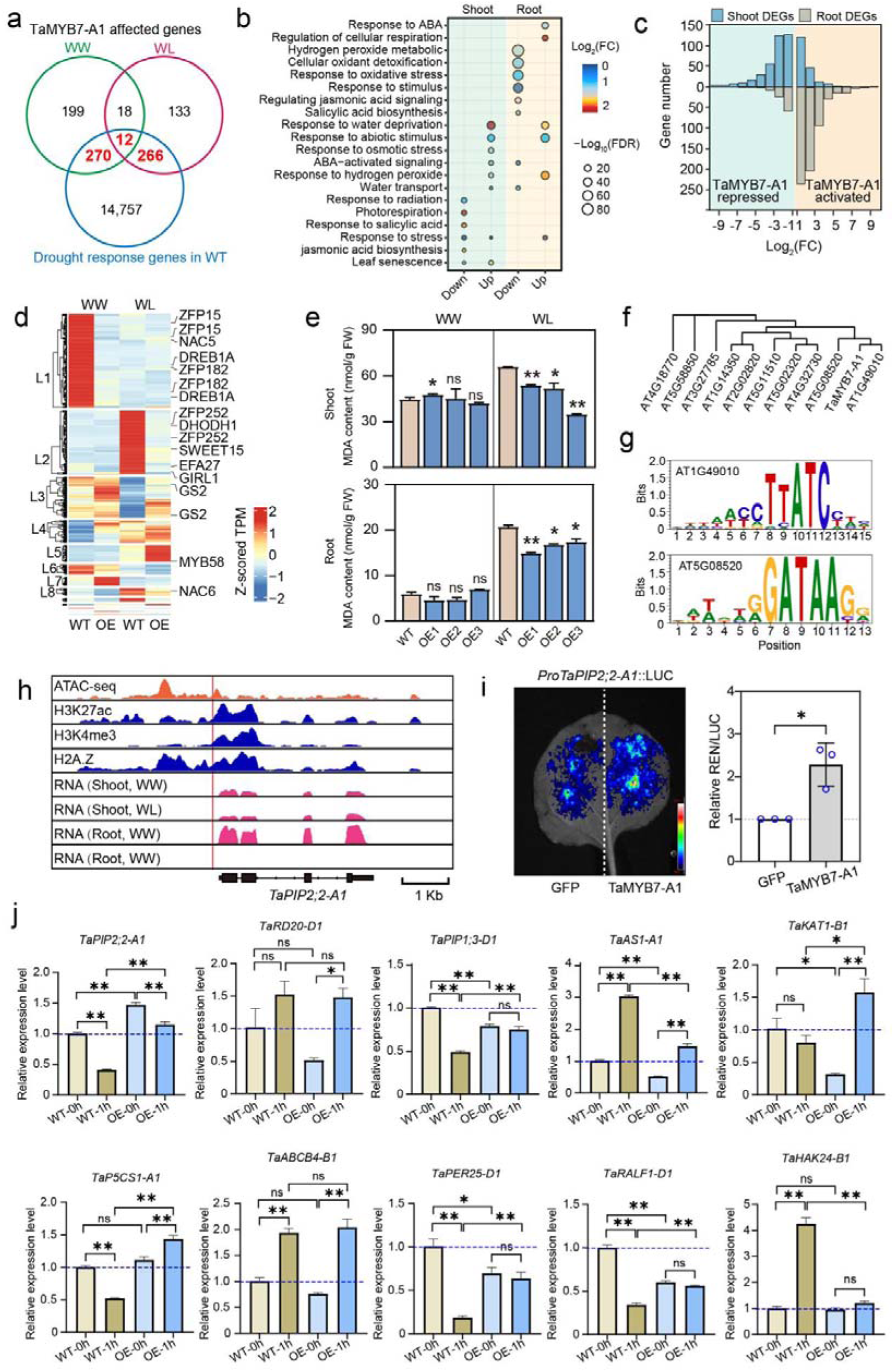
RNA-seq identifying potential downstream direct targets of TaMYB7-A1. **(a)** Venn diagram displaying the overlap of TaMYB7-A1 affected genes under WW (green) and WL (red) conditions and drought-response genes (blue) in shoot. The TaMYB7-A1 affected drought response genes (DRGs) were marked in red. DEGs were identified with Log2 (FoldChange)>1 and *P*.adjust<0.05. **(b)** Enriched Gene Ontology terms for drought response genes DRGs in shoot and root. The size of dots indicates gene ratio enriched in each term, with color representing the *P*.adjust value, which refers to the Benjamini-Hochberg procedure adjusted *p*-values of Hypergeometric test. **(c)** Distribution of the TaMYB7-A1 -activated (yellow shade) and -repressed (blue shade) genes in shoot (blue bars) and root (gray bars) across different -log_2_FC values. The foldchange were calculated using the mean values of TPM_OE_/TPM_Filder_ in three OE lines. **(d)** Heatmap showing the k-means clustering of TaMYB7-A1 affected DRGs in shoot, with reported drought-resistance genes labeled. The color intensity from blue to red represents the z-scored TPM values ranging from -2 to 2. **(e)** Comparison of the MDA content between seedlings of Fielder and OE lines under WW and WL conditions. Data are means ± S.D. (*n* = 4 independent biological replicates). Student’s *t* test were used to determine the statistical significance between each OE line and Fielder. ns, no significance; *, *P* < 0.05; **, *P* < 0.01. (**f, g**) Neighbor-joining phylogenetic tree of TaMYB7-A (e) and the binding motif of its closest Arabidopsis orthologs (f). Arabidopsis orthologs AT1G49010 and AT5G08520 were served as alternatives of *TaMYB7-A1*, and their binding motif for screening binding motifs of TaMYB7-A1. The phylogeny testing applied the bootstrap method with 1,000 replicates. **(h)** IGV showing the RNA-seq, ATAC-seq, and each histone modification across *TaPIP2;2-A1*. Red vertical line indicates the predicted binding motif of TaMYB7-A1. **(i)** Dual-luciferase (LUC) reporter assays showing the transcriptional activation of TaMYB7-A1 on *TaPIP2;2-B1*. The relative value of LUC/REN (Renilla) was normalized with value in green fluorescent protein (GFP) set as 1. Error bars show ± S.D. of biological replicates (*n* = 3). Student’s *t*-test was used for the statistical significance. *, *P* < 0.05. **(j)** Expression levels of several potential downstream direct targets of TaMYB7-A1 in root of Fielder and OE1 under 12% PEG-simulated drought stress. RT-qPCR was used to determine their expression patterns, with *TaTublin* as the internal control. The expression levels were normalized with WT at 0 hour set to 1.0. Data are means ± S.D. of three independent biological replicates. Student’s *t* test were used to determine the statistical significance. no significant difference; *, *P* < 0.05; **, *P* < 0.01. h, hour.

## Reference

1. Fao et al. The State of Food Security and Nutrition in the World 2020. Transforming food systems for affordable healthy diets. IEEE Journal of Selected Topics in Applied Earth Observations and Remote Sensing (2020).

2. Hussain, J., Khaliq, T., Ahmad, A., Akhter, J. & Asseng, S. Wheat Responses to Climate Change and Its Adaptations: A Focus on Arid and Semi-arid Environment. Int J Environ Res 12, 117–126 (2018).

3. Trnka, M. et al. Mitigation efforts will not fully alleviate the increase in water scarcity occurrence probability in wheat-producing areas. Science Advances (2019) doi:10.1126/sciadv.aau2406.

4. Rezaei, E. E. et al. Climate change impacts on crop yields. Nature Reviews Earth & Environment 4, 831–846 (2023).

5. Leghari, S. J. et al. What should we do for water security? A technical review on more yield per water drop. Journal of Environmental Management 370, 122832 (2024).

6. Bapela, T., Shimelis, H., Tsilo, T. J. & Mathew, I. Genetic Improvement of Wheat for Drought Tolerance: Progress, Challenges and Opportunities. Plants 11, 1331 (2022).

7. Ali, O. A. M. Wheat Responses and Tolerance to Drought Stress. in Wheat Production in Changing Environments: Responses, Adaptation and Tolerance (eds. Hasanuzzaman, M., Nahar, K. & Hossain, Md. A.) 129–138 (Springer, Singapore, 2019). doi:10.1007/978-981-13-6883-7_5.

8. Gupta, A., Rico-Medina, A. & Caño-Delgado, A. I. The physiology of plant responses to drought. Science 368, 266–269 (2020).

9. Abid, M. et al. Physiological and biochemical changes during drought and recovery periods at tillering and jointing stages in wheat (Triticum aestivum L.). Scientific reports 8, 4615 (2018).

10. Tomar, R. S. S. et al. Molecular and Morpho-Agronomical Characterization of Root Architecture at Seedling and Reproductive Stages for Drought Tolerance in Wheat. PLoS ONE 11, e0156528 (2016).

11. Cardoso, A. A., Gori, A., Da-Silva, C. J. & Brunetti, C. Abscisic acid biosynthesis and signaling in plants: Key targets to improve water use efficiency and drought tolerance. Applied Sciences 10, 6322 (2020).

12. Blatt, M. R., Jezek, M., Lew, V. L. & Hills, A. What can mechanistic models tell us about guard cells, photosynthesis, and water use efficiency? Trends in Plant Science 27, 166–179 (2022).

13. Chen, J. et al. TaNAC48 positively regulates drought tolerance and ABA responses in wheat (Triticum aestivum L.). The Crop Journal (2020) doi:10.1016/j.cj.2020.09.010.

14. Maghsoudi, K., Emam, Y., Niazi, A., Pessarakli, M. & Arvin, M. J. P5CS expression level and proline accumulation in the sensitive and tolerant wheat cultivars under control and drought stress conditions in the presence/absence of silicon and salicylic acid. Journal of Plant Interactions 13, 461–471 (2018).

15. Mei, F. et al. A gain-of-function allele of a DREB transcription factor gene ameliorates drought tolerance in wheat. The Plant Cell koac248 (2022) doi:10.1093/plcell/koac248.

16. Wang, J., et al. *DIW1* encoding a clade I PP2C phosphatase negatively regulates drought tolerance by deLphosphorylating TaSnRK1.1 in wheat. JIPB 65, 1918–1936 (2023).

17. Mao, H. et al. The wheat ABA receptor gene TaPYL1-1B contributes to drought tolerance and grain yield by increasing water-use efficiency. Plant Biotechnology Journal n/a,.

18. Zhang, Y. et al. TaPYL4, an ABA receptor gene of wheat, positively regulates plant drought adaptation through modulating the osmotic stress-associated processes. BMC Plant Biol 22, 423 (2022).

19. Zhang, Y. et al. Wheat TaPYL9 Linvolved signalling pathway impacts plant drought response through regulating distinct osmotic stressLassociated physiological indices. Plant Biotechnology Journal pbi.14501 (2024) doi:10.1111/pbi.14501.

20. Impa, S. M., Nadaradjan, S. & Jagadish, S. V. K. Drought Stress Induced Reactive Oxygen Species and Anti-oxidants in Plants. in Abiotic Stress Responses in Plants (eds. Ahmad, P. & Prasad, M. N. V.) 131–147 (Springer New York, New York, NY, 2012). doi:10.1007/978-1-4614-0634-1_7.

21. Hasanuzzaman, M., Nahar, K., Gill, S. S. & Fujita, M. Drought Stress Responses in Plants, Oxidative Stress, and Antioxidant Defense. in Climate Change and Plant Abiotic Stress Tolerance (eds. Tuteja, N. & Gill, S. S.) 209–250 (Wiley, 2013). doi:10.1002/9783527675265.ch09.

22. Caverzan, A., Casassola, A. & Brammer, S. P. Antioxidant responses of wheat plants under stress. Genetics and molecular biology 39, 1–6 (2016).

23. Singh, M., Kumar, J., Singh, S., Singh, V. P. & Prasad, S. M. Roles of osmoprotectants in improving salinity and drought tolerance in plants: a review. Rev Environ Sci Biotechnol 14, 407–426 (2015).

24. Shi, X. et al. The mechanism of Ca2+ signal transduction in plants responding to abiotic stresses. Environmental and Experimental Botany 105514 (2023).

25. Khatiwada, A., Neupane, I., Sharma, B., Bhetwal, N. & Pandey, B. Effects of Drought Stress on Yield and Yield Attributing Characters of Wheat: A Review. agw 8, 118–124 (2020).

26. Leakey, A. D. B. et al. Water Use Efficiency as a Constraint and Target for Improving the Resilience and Productivity of C _3_ and C _4_ Crops. Annu. Rev. Plant Biol. 70, 781–808 (2019).

27. Tardieu, F. Different avenues for progress apply to drought tolerance, water use efficiency and yield in dry areas. Current Opinion in Biotechnology 73, 128–134 (2022).

28. Yang, Z. & Qin, F. The battle of crops against drought: Genetic dissection and improvement. JIPB 65, 496–525 (2023).

29. Gupta, P. K., Balyan, H. S. & Gahlaut, V. QTL analysis for drought tolerance in wheat: present status and future possibilities. Agronomy 7, 5 (2017).

30. Mao, H. et al. Variation in cis-regulation of a NAC transcription factor contributes to drought tolerance in wheat. Molecular Plant 15, 276–292 (2022).

31. Zhang, L. et al. A wheat R2R3-MYB gene, TaMYB30-B, improves drought stress tolerance in transgenic Arabidopsis. Journal of experimental botany 63, 5873–5885 (2012).

32. Bu, Y. et al. The transcription factor TabZIP156 acts as a positive regulator in response to drought tolerance in Arabidopsis and wheat (Triticum aestivum L.). Plant Physiology and Biochemistry 216, 109086 (2024).

33. Ge, M. et al. TaWRKY31, a novel WRKY transcription factor in wheat, participates in regulation of plant drought stress tolerance. BMC Plant Biol 24, 27 (2024).

34. Wang, D. et al. TabHLH27 orchestrates root growth and drought tolerance to enhance water use efficiency in wheat. Journal of Integrative Plant Biology 66, 1295–1312 (2024).

35. Placido, D. F. et al. The *LATERAL ROOT DENSITY* gene regulates root growth during water stress in wheat. Plant Biotechnology Journal 18, 1955–1968 (2020).

36. Ma, J. et al. TaSINA2B, interacting with TaSINA1D, positively regulates drought tolerance and root growth in wheat (*Triticum aestivum* L.). Plant Cell & Environment 46, 3760–3774 (2023).

37. Li, J. et al. The potassium transporter TaNHX2 interacts with TaGAD1 to promote drought tolerance via modulating stomatal aperture in wheat. Sci. Adv. 10, eadk4027 (2024).

38. He, J., et al. *ECERIFERUM1-6A* is required for the synthesis of cuticular wax alkanes and promotes drought tolerance in wheat. Plant Physiology 190, 1640–1657 (2022).

39. Tian, G. et al. Allelic variation of TaWD40–4B.1 contributes to drought tolerance by modulating catalase activity in wheat. Nat Commun 14, 1200 (2023).

40. Zhang, L. et al. The wheat VQ motif-containing protein TaVQ4-D positively regulates drought tolerance in transgenic plants. Journal of Experimental Botany 74, 5591–5605 (2023).

41. Mwadzingeni, L., Shimelis, H., Dube, E., Laing, M. D. & Tsilo, T. J. Breeding wheat for drought tolerance: Progress and technologies. Journal of Integrative Agriculture 15, 935–943 (2016).

42. Li, Z. et al. Combined GWAS and eQTL analysis uncovers a genetic regulatory network orchestrating the initiation of secondary cell wall development in cotton. New Phytologist doi:10.1111/nph.16468.

43. Han, X. et al. Integration of eQTL Analysis and GWAS Highlights Regulation Networks in Cotton under Stress Condition. Int J Mol Sci 23, 7564 (2022).

44. Lin, X. et al. Systematic identification of wheat spike developmental regulators by integrated multi-omics, transcriptional network, GWAS, and genetic analyses. Molecular Plant 17, 438–459 (2024).

45. Yuan, X. et al. Integrative omics analysis elucidates the genetic basis underlying seed weight and oil content in soybean. The Plant Cell 36, 2160–2175 (2024).

46. Zhao, L. et al. Deciphering the Transcriptional Regulatory Network Governing Starch and Storage Protein Biosynthesis in Wheat for Breeding Improvement. Advanced Science n/a, 2401383.

47. Liu, S. et al. Mapping regulatory variants controlling gene expression in drought response and tolerance in maize. Genome Biol 21, 163 (2020).

48. Ma, L. et al. GWAS and WGCNA uncover hub genes controlling salt tolerance in maize (Zea mays L.) seedlings. Theor Appl Genet 134, 3305–3318 (2021).

49. Yang, G. et al. Combined GWAS and eGWAS reveals the genetic basis underlying drought tolerance in emmer wheat (*Triticum turgidum* L.). New Phytologist 242, 2115–2131 (2024).

50. Wang, D. et al. Boosting wheat functional genomics via an indexed EMS mutant library of KN9204. Plant Commun 4, 100593 (2023).

51. Sandhu, N. et al. Rice root architectural plasticity traits and genetic regions for adaptability to variable cultivation and stress conditions. Plant physiology 171, 2562–2576 (2016).

52. Qiao, Y. et al. Development and application of a relative soil water content – transpiration efficiency curve for screening high water use efficiency wheat cultivars. Frontiers in Plant Science 13, (2022).

53. Sun, C. et al. The Wheat 660K SNP array demonstrates great potential for marker-assisted selection in polyploid wheat. Plant Biotechnology Journal 18, 1354–1360 (2020).

54. Li, A. L. et al. Wheat breeding history reveals synergistic selection of pleiotropic genomic sites for plant architecture and grain yield. Mol Plant 15, 504–519 (2022).

55. Pang, Y. et al. High-Resolution Genome-wide Association Study Identifies Genomic Regions and Candidate Genes for Important Agronomic Traits in Wheat. Molecular Plant 13, 1311–1327 (2020).

56. Jin, M. et al. Maize pan-transcriptome provides novel insights into genome complexity and quantitative trait variation. Sci Rep 6, 18936 (2016).

57. Bhatti, A., Shah, F. S., Azhar, J., Ahmad, S. & John, P. Chapter 18 - Pan-transcriptomics and its applications. in Pan-genomics: Applications, Challenges, and Future Prospects (eds. Barh, D., Soares, S., Tiwari, S. & Azevedo, V.) 343–356 (Academic Press, 2020). doi:10.1016/B978-0-12-817076-2.00018-4.

58. Li, J. et al. The potassium transporter TaNHX2 interacts with TaGAD1 to promote drought tolerance via modulating stomatal aperture in wheat. Science Advances 10, eadk4027 (2024).

59. Du, L. et al. TaERF87 and TaAKS1 synergistically regulate TaP5CS1 / TaP5CR1 Lmediated proline biosynthesis to enhance drought tolerance in wheat. New Phytologist 237, 232–250 (2023).

60. Ramírez-González, R. H. et al. The transcriptional landscape of polyploid wheat. Science 361, eaar6089 (2018).

61. Wang, M. et al. An atlas of wheat epigenetic regulatory elements reveals subgenome divergence in the regulation of development and stress responses. The Plant Cell 33, 865–881 (2021).

62. He, F. et al. Genomic variants affecting homoeologous gene expression dosage contribute to agronomic trait variation in allopolyploid wheat. Nat Commun 13, 826 (2022).

63. Deng, P. et al. Population-level transcriptomes reveal gene expression and splicing underlying cadmium accumulation in barley. The Plant Journal 112, 847–859 (2022).

64. Cookson, W., Liang, L., Abecasis, G., Moffatt, M. & Lathrop, M. Mapping complex disease traits with global gene expression. Nat Rev Genet 10, 184–194 (2009).

65. Li, Z. et al. Combined GWAS and eQTL analysis uncovers a genetic regulatory network orchestrating the initiation of secondary cell wall development in cotton. New Phytologist 226, 1738–1752 (2020).

66. Vasudevan, A., Lévesque-Lemay, M., Edwards, T. & Cloutier, S. Global transcriptome analysis of allopolyploidization reveals large-scale repression of the D-subgenome in synthetic hexaploid wheat. Communications Biology 6, 426 (2023).

67. Zhao, G. et al. The Aegilops tauschii genome reveals multiple impacts of transposons. Nature Plants 3, 946–955 (2017).

68. Tang, Y. et al. Overexpression of a MYB family gene, OsMYB6, increases drought and salinity stress tolerance in transgenic rice. Frontiers in plant science 10, 168 (2019).

69. Yuan, F. et al. OSCA1 mediates osmotic-stress-evoked Ca2+ increases vital for osmosensing in Arabidopsis. Nature 514, 367–371 (2014).

70. Van Norman, J. M., Frederick, R. L. & Sieburth, L. E. BYPASS1 negatively regulates a root-derived signal that controls plant architecture. Current Biology 14, 1739–1746 (2004).

71. Takahashi, Y. et al. bHLH Transcription Factors That Facilitate K+ Uptake During Stomatal Opening Are Repressed by Abscisic Acid Through Phosphorylation. Science Signaling 6, ra48–ra48 (2013).

72. Song, C.-P. et al. Role of an Arabidopsis AP2/EREBP-type transcriptional repressor in abscisic acid and drought stress responses. The Plant Cell 17, 2384–2396 (2005).

73. Wang, H. et al. Natural variations of TaMYB7–A1 regulate PHS resistance and specify wheat geographic adaptation. 2024.11.15.623828 Preprint at 10.1101/2024.11.15.623828 (2024).

74. Tournaire-Roux, C. et al. Cytosolic pH regulates root water transport during anoxic stress through gating of aquaporins. Nature 425, 393–397 (2003).

75. Lian, H.-L. et al. The Role of Aquaporin RWC3 in Drought Avoidance in Rice. Plant and Cell Physiology 45, 481–489 (2004).

76. Park, S.-I., Kim, J.-J., Shin, S.-Y., Kim, Y.-S. & Yoon, H.-S. ASR Enhances Environmental Stress Tolerance and Improves Grain Yield by Modulating Stomatal Closure in Rice. Front. Plant Sci. 10, 1752 (2020).

77. James, D. et al. Concurrent Overexpression of OsGS1;1 and OsGS2 Genes in Transgenic Rice (Oryza sativa L.): Impact on Tolerance to Abiotic Stresses. Front. Plant Sci. 9, (2018).

78. Hirano, K. et al. Identification of Transcription Factors Involved in Rice Secondary Cell Wall Formation. Plant and Cell Physiology 54, 1791–1802 (2013).

79. Zhang, H. et al. The C2H2-type Zinc Finger Protein ZFP182 is Involved in Abscisic Acid-Induced Antioxidant Defense in Rice. Journal of Integrative Plant Biology 54, 500–510 (2012).

80. Aubert, Y. et al. RD20, a Stress-Inducible Caleosin, Participates in Stomatal Control, Transpiration and Drought Tolerance in Arabidopsis thaliana. Plant and Cell Physiology 51, 1975–1987 (2010).

81. Bandurska, H. et al. Regulation of proline biosynthesis and resistance to drought stress in two barley (Hordeum vulgare L.) genotypes of different origin. Plant Physiology and Biochemistry 118, 427–437 (2017).

82. Crick, F. Central dogma of molecular biology. Nature 227, 561–563 (1970).

83. Korte, A. & Farlow, A. The advantages and limitations of trait analysis with GWAS: a review. Plant Methods 9, 29 (2013).

84. Hu, H. & Xiong, L. Genetic engineering and breeding of drought-resistant crops. Annual review of plant biology 65, 715–741 (2014).

85. Huang, X. & Han, B. Natural Variations and Genome-Wide Association Studies in Crop Plants. Annu. Rev. Plant Biol. 65, 531–551 (2014).

86. Morsy, A., El-Sherif, N., Morsy, A. & El-Sherif, N. Integrating Omics Approaches for Abiotic Stress Tolerance in Plants. in Abiotic Stress in Crop Plants - Ecophysiological Responses and Molecular Approaches (IntechOpen, 2024). doi:10.5772/intechopen.114121.

87. Yang, Y. et al. Applications of Multi-Omics Technologies for Crop Improvement. Front. Plant Sci. 12, (2021).

88. Han, L. et al. A multi-omics integrative network map of maize. Nat Genet 55, 144–153 (2023).

89. Rivera, H. E. et al. A framework for understanding gene expression plasticity and its influence on stress tolerance. Molecular Ecology 30, 1381–1397 (2021).

90. López-Maury, L., Marguerat, S. & Bähler, J. Tuning gene expression to changing environments: from rapid responses to evolutionary adaptation. Nature Reviews Genetics 9, 583–593 (2008).

91. Van de Peer, Y., Mizrachi, E. & Marchal, K. The evolutionary significance of polyploidy. Nature Reviews Genetics 18, 411–424 (2017).

92. Sharma, N., Raman, H., Wheeler, D., Kalenahalli, Y. & Sharma, R. Data-driven approaches to improve water-use efficiency and drought resistance in crop plants. Plant Science 336, 111852 (2023).

93. Hafeez, A. et al. Breeding for water-use efficiency in wheat: progress, challenges and prospects. Mol Biol Rep 51, 429 (2024).

94. Condon, A. G., Richards, R., Rebetzke, G. & Farquhar, G. Breeding for high water-use efficiency. Journal of experimental botany 55, 2447–2460 (2004).

95. Zhou, X. Genome-wide efficient mixed-model analysis for association studies. Nature Genetics 44, (2012).

96. Shabalin, A. A. Matrix eQTL: ultra fast eQTL analysis via large matrix operations. Bioinformatics 28, 1353–1358 (2012).

97. Li, M.-X., Yeung, J. M. Y., Cherny, S. S. & Sham, P. C. Evaluating the effective numbers of independent tests and significant p-value thresholds in commercial genotyping arrays and public imputation reference datasets. Hum Genet 131, 747–756 (2012).

98. Zhang, B. & Horvath, S. A General Framework for Weighted Gene Co-Expression Network Analysis. Statistical Applications in Genetics and Molecular Biology 4, (2005).

99. Zhu, Z. et al. Integration of summary data from GWAS and eQTL studies predicts complex trait gene targets. Nat Genet 48, 481–487 (2016).

100. You, J. et al. Regulatory controls of duplicated gene expression during fiber development in allotetraploid cotton. Nat Genet 55, 1987–1997 (2023).

101. Liu, Z. et al. Temporal transcriptome profiling reveals expression partitioning of homeologous genes contributing to heat and drought acclimation in wheat (Triticum aestivum L.). BMC Plant Biol 15, 152 (2015).

